# Integrated single-cell profiling of RNA and DNA interactomes reveals targetable chromatin architectures in cancer

**DOI:** 10.64898/2026.08.12.744478

**Authors:** Evgeny Deforzh, Yanhong Zhang, Hayk Mnatsakanyan, Alden John, Amelie Kinsey, Zelong Zheng, Abdellatif El Khayari, Christian E. Badr, Anna M. Krichevsky

## Abstract

The human genome is pervasively transcribed into protein-coding and regulatory non-coding RNAs whose functions are coordinated within higher-order nuclear architectures. However, direct mapping of RNA–RNA and DNA–DNA interaction networks in complex human tissues at single-cell resolution has remained a major challenge. Here, we present SCIENCE-seq, a multimodal single-cell interactomics platform enabling simultaneous detection of RNA–RNA interactions and chromatin DNA–DNA contacts within individual cells. Applied to primary glioma specimens, SCIENCE-seq reveals cancer-specific interactions organized by lncRNAs and centered on key oncogenic drivers, including EGFR, hTERT, SOX2, CDK6, CDC42, CD47, and HOX loci. These datasets uncover previously unrecognized molecular relationships between premature lncRNAs and pre-mRNAs that regulate transcription, alternative splicing, and polyadenylation. Notably, we identify a glioma-specific trans-chromosomal HOX hub driven by five interacting lncRNAs. Targeting specific RNA and DNA interactions using steric antisense oligonucleotides (ASOs) or CRISPRi selectively suppresses oncogenic programs in malignant cells while sparing normal tissue, establishing chromatin interactomes as actionable therapeutic targets.

---

The establishment of cell-type-specific gene expression programs is fundamental to development and is frequently disrupted in complex diseases, including cancer^1,2^. This regulation extends beyond linear DNA sequence and is governed by a dynamic three-dimensional nuclear architecture^3,4^, in which distal genomic elements-including enhancers, silencers, and non-coding RNA loci-communicate through spatial proximity.

Emerging evidence indicates that this architecture is shaped not only by DNA–DNA contacts but also by extensive RNA–RNA interactions that contribute to nuclear organization and gene regulation^5,6^. These multimolecular networks are thought to coordinate multiple stages of RNA biogenesis, including transcription initiation^7^, alternative splicing^8^, and potentially alternative polyadenylation. In cancer, dysregulation of these spatial networks enables the formation of regulatory “interactomic hubs” that sustain oncogenic gene expression programs^9,10^.

Despite their importance, these networks remain poorly characterized. Existing approaches typically measure DNA–DNA interactions (Hi-C^4^, ChIA-PET^11^, SPRITE^12^) or RNA–RNA interactions (PARIS^13^, SPLASH^14^, RIC-seq^10^) independently, resulting in fragmented views of nuclear organization. Moreover, most methods rely on bulk measurements, obscuring cell-type-specific heterogeneity. High-throughput imaging-based approaches such as seqFISH+^15^ and MERFISH^16^ provide single cell spatial resolution but do not directly resolve molecular interactions. Recent single-cell methods such as scSPRITE^8^ detect multi-molecular DNA complexes but lack the resolution to define discrete pairwise interactions.

Here, we address these limitations by developing a multimodal single-cell interactomics platform that simultaneously captures RNA–RNA and DNA–DNA interactions within individual cells. By combining high-resolution proximity ligation with single-cell barcoding, this approach enables direct mapping of interacting RNA segments alongside their corresponding chromatin contacts.

Applying this platform to primary high-grade glioma tissues, we uncover previously unrecognized regulatory architectures organized by lncRNAs around key oncogenic drivers, including SOX2, EGFR, CDK6, hTERT, CDC42, CD47, and HOX loci. These data reveal that lncRNAs form cell-type-specific interactions with pre-mRNAs that control multiple layers of gene regulation, including transcription, alternative splicing, and polyadenylation.

Importantly, we demonstrate that these RNA-centered interaction hubs can be selectively disrupted using steric antisense oligonucleotides (ASOs), enabling targeted inhibition of oncogenic programs in glioma cells while sparing normal brain cell populations. Together, our findings establish RNA and DNA interactomes as a previously unrecognized and therapeutically tractable layer of gene regulation in complex human tissues.

## SCIENCE-seq enables multimodal single-cell mapping of RNA–RNA and DNA–DNA interactions

To simultaneously interrogate RNA–RNA and DNA–DNA interactions within individual cells, we developed **Single-Cell Integrated Evaluation of Nucleic Contact Events (SCIENCE-seq)**, a multimodal interactomics platform (**Fig. 1A**). SCIENCE-seq integrates proximity ligation–based detection of molecular interactions with single-cell barcoding, enabling parallel reconstruction of RNA and chromatin interaction networks.

**Figure 1.**
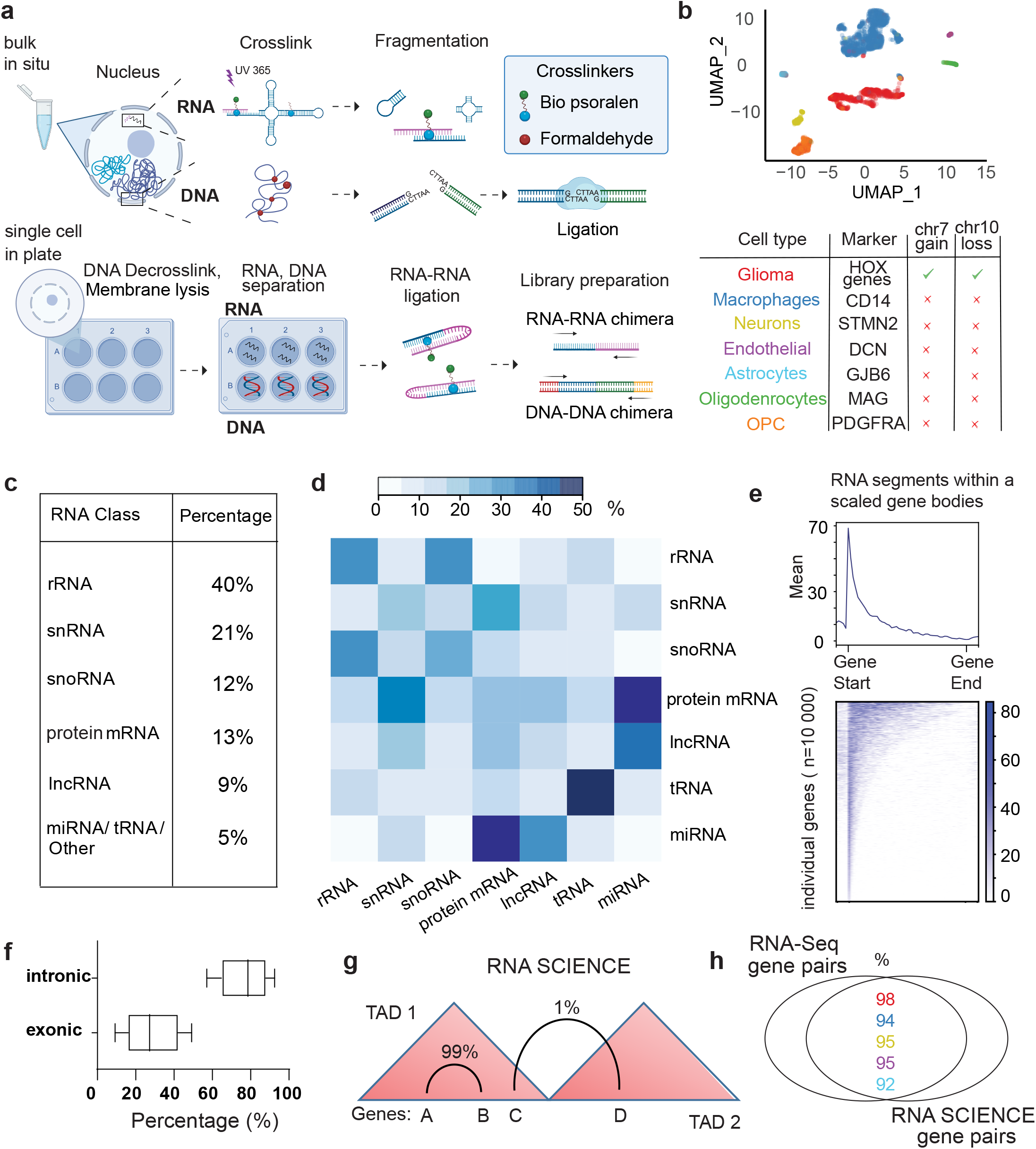
Combined single cell RNA-RNA and DNA-DNA interactomics (SCIENCE-seq) a. Diagram of the combined single cell RNA-RNA and DNA-DNA interactomic technique (SCIENCE-seq). b. Single-cell interactomic landscape of primary glioma. UMAP visualization of ∼3,500 cells (dots) from 3 patients, with clusters resolved exclusively by RNA-RNA interactomic profiles. c. Distribution of the RNA interactome by RNA biotype (percentage). d. Normalized RNA-RNA interactomic matrix. Heatmap showing the row-normalized distribution of RNA-RNA interactions in percentage (color bar on top), highlighting the preferential target classes for each specific RNA biotype, independent of total interaction counts. e. Enrichment of RNA segments at the 5′ termini. RNA segments, visualized within scaled gene bodies (n=10,000) as average signal (top line plot) or individual raw read counts (bottom heatmap) are preferentially localized at the beginning of the gene body, primarily originating from the first exon or the following intron. f. Distribution of intronic versus exonic interacting segments. Intronic and exonic segments were quantified within protein-coding genes and lncRNAs for each cell (n=3500) and visualized as boxplots. g. Positioning of interacting intronic RNA pairs relative to TAD boundaries. Integration of Hi-C and RNA SCIENCE datasets in glioma cells reveals that the vast majority (99%) of intronic associations are spatially constrained within the same Topologically Associating Domains (TADs). h. High concordance between RNA interactomes and transcriptional co-expression. Percentage of SCIENCE-seq gene pairs matching intra-TAD gene pairs with the highest expression correlation (Pearson coefficients) across five cell types.

We applied SCIENCE-seq to primary glioblastoma (GBM) specimens from three patients. Single-cell suspensions prepared from frozen tumors were subjected to independent crosslinking of RNA– RNA and DNA–DNA interactions, followed by fragmentation and proximity ligation. After redistribution into 96-well plates, crosslinks were reversed and RNA and DNA fractions were processed separately for library preparation (see Methods).

Single-cell transcriptomes derived from RNA SCIENCE data were used to cluster cells by UMAP (**Fig. 1B**). Glioma cells were distinguished from non-malignant populations based on canonical marker expression and chromosome 7/10 copy number variation. Consistent with prior studies, tumors comprised heterogeneous populations including glioma cells, macrophages, OPCs, and additional neural lineages. Notably, expression of HOX genes alone robustly separated glioma cells from normal brain cell types, suggesting a distinct transcriptional program associated with malignancy.

Across individual cells, SCIENCE-seq detected approximately ∼7,000 expressed genes and ∼2 million RNA–RNA interactions (∼80,000 unique pairs of interacting segments), with interaction lengths averaging ∼40 nucleotides and ∼80% complementarity. To validate interaction specificity, we examined canonical interactions between 45S pre-rRNA and C/D box snoRNAs (SNORDs), which were accurately recovered through their known guide sequences (**Supplementary Fig. 1A, B**). In addition, we identified a broad repertoire of interactions involving snRNAs, SNORDs, and SNORAs, including previously uncharacterized associations **(Supplementary Fig. 1B, Supplementary Table 1)**.

Globally, RNA SCIENCE captured both inter- and intra-molecular interactions across major RNA classes (**Fig. 1C, D**), including known interactions (e.g., rRNA–snoRNA, miRNA–mRNA) and less characterized ones (e.g., miRNA–lncRNA, lncRNA–lncRNA). Interaction sites were strongly enriched within the first ∼1 kb downstream of transcription start sites (TSSs) (**Fig. 1E**), corresponding to first exons or proximal intronic regions in protein-coding and lncRNA transcripts (**Fig. 1F**).

Integration with public RNA-seq and Hi-C datasets revealed that interacting RNA pairs are highly enriched among co-expressed genes within the same topologically associating domains (TADs) (**Fig. 1G**). Notably, 92–98% concordance was observed between gene pairs with the strongest expression correlations and those identified as interacting by RNA SCIENCE (**Fig. 1H**), suggesting that RNA–RNA interactions reflect coordinated transcriptional regulation within chromatin domains. For simplicity, overlapping RNA interaction segments were merged into common interaction regions (illustrated in **Supplemental Fig. 1C**) in all subsequent analyses and figures. Consistently, DNA SCIENCE identified promoter-proximal interactions among regulatory elements within the same TADs, indicating that RNA–RNA and DNA–DNA contacts converge on shared regulatory architectures (**Supplementary Fig. 1D**).

### SCIENCE-seq reveals cell-type specific interactions between nascent lncRNA- and oncogenic drivers

We next investigated whether multimodal interaction networks organize gene regulation in a cell-type-specific manner within tumors. SCIENCE-seq revealed promoter-promoter and single-nucleotide resolution RNA–RNA interactions linking cell-type-specific nascent lncRNAs to key oncogenic drivers, including *SOX2*, *hTERT, CDK6*, and *EGFR* (**Fig. 2A, B; Supplemental Figure 2A-C).** Notably, some individual oncogenes exhibited mutually exclusive interactions with different lncRNAs across cell types, suggesting that lncRNA identity encodes cell-type-specific regulatory states (**Fig. 2B**).

**Figure 2.**
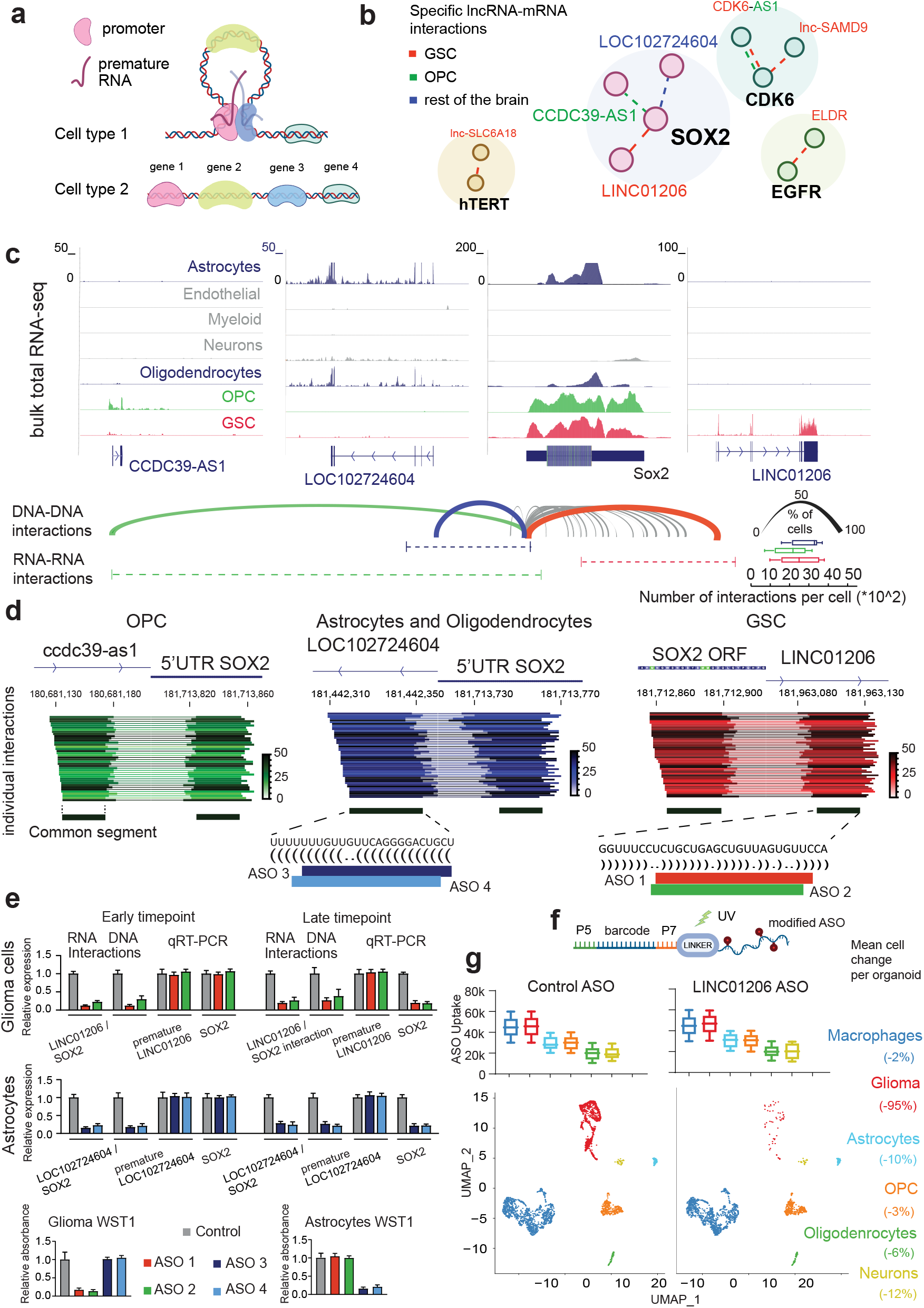
Integrated RNA and DNA interactomics reveals a comprehensive landscape of interactions between disease-specific lncRNA and critical glioma drivers. a. Cell type-specific premature RNA-RNA and promoter DNA-DNA interactions facilitate the spatial proximity of distal genes. b. Cell type-specific RNA interactomic networks. RNA-RNA interactions (dotted lines) between lncRNAs and key protein-coding glioma drivers are colored by cell type. c. Integrated RNA and DNA interactomics identify cell type-specific contacts between SOX2 and neighboring lncRNAs. Total Bulk RNA-Seq in different cell types (top) shows cell type-specific lncRNA expression. SCIENCE-Seq shows cell type-specific DNA and RNA contacts between common interacting segments (bottom). Arc width (bottom, right) represents the percentage of cells with a corresponding DNA-DNA interaction. Boxplots (bottom right) quantify the frequency of RNA-RNA interactions per cell. d. Mapping of individual interacting RNA segments for therapeutic targeting. Intragenic positioning and interaction counts for SOX2-lncRNA segments (from Fig. 2C) are shown across OPCs, Astrocytes / Oligodendrocytes and Glioma stem cells. The common segments of LINC01206 or LOC102724604 are highlighted in black, with their primary sequences and predicted base-pairing complementarity to SOX2 detailed below. Antisense Oligonucleotides (ASO) designed to target those common segments within corresponding lncRNAs are shown as colored lines at the bottom. e. Targeted disruption of the cell-specific RNA and DNA interactomes with ASOs. Treatment of glioma stem cells or astrocytes with two ASOs disrupts cell type-specific premature RNA interactions (SPLASH; n=3, mean with SD) and DNA loops (3C-qPCR; n=3, mean with SD) between SOX2 and corresponding cell-type specific lncRNA, leading to decreased SOX2 expression (qRT-PCR, mean with SD, n=3) and cell death (WST1, mean with SD, n=3). Early-timepoint analysis reveals that ASO treatment successfully dissociates RNA and DNA interactions prior to any measurable reduction in steady-state transcript levels. f. Design of the Barcoded ASO. Barcode sequence made of 20 random nucleotides flanked by Illumina-compatible overhangs, was conjugated to the ASO via a UV-cleavable linker to enable controlled release and subsequent sequencing (see Methods for details). g. LINC01206 ASO treatment induces selective glioma cell death. Organoids (n=2) were treated with Control ASO or LINC01206 ASO (ASO1 from Figure 2F). ASO uptake (upper panel) was measured for each individual cell, grouped by cell type and visualized with boxplots. Synchronized UMAP plots (lower panel) display cell clusters following treatment with Control or LINC01206 ASO. Mean cell count changes per organoid (Right). Every dot is an individual cell.

Focusing on the stemness regulator SOX2, we observed that its expression varies across tumor cell populations, with highest levels in glioma stem cells (GSCs) and oligodendrocyte precursor cells (OPCs), moderate expression in astrocytes and oligodendrocytes, and minimal or absent expression in neurons, endothelial, and myeloid populations. Three neighboring lncRNAs-LINC01206, CCDC39-AS1, and LOC102724604 - exhibited mutually exclusive expression patterns across these same cell populations: LINC01206 was selectively expressed in GSCs, CCDC39-AS1 in OPCs, and LOC102724604 in astrocytes and oligodendrocytes (**Fig. 2C**).

DNA SCIENCE analysis revealed that the SOX2 genomic locus forms cell-type-specific chromatin contacts with each of these lncRNA loci (**Fig. 2C**, bottom arcs). Concordantly, RNA SCIENCE identified direct and mutually exclusive interactions between SOX2 mRNA and these lncRNAs across the corresponding cell types: LINC01206 interacted with SOX2 specifically in glioma cells, CCDC39-AS1 in OPCs, and LOC102724604 in astrocyte/oligodendrocyte populations (**Fig. 2C**, common interacting segments, bottom lines). Common interacting regions across cell types were composed of similarly sized individual segments with modestly shifted genomic boundaries and at least 80% complementarity **(Fig. 2D).**

To determine whether these interactions are functionally required for SOX2 regulation, we used steric-blocking ASOs targeting the common SOX2-interacting regions of LINC01206 (in glioma cells) and LOC102724604 (in astrocytes). ASO-mediated disruption reduced both RNA–RNA and DNA–DNA contacts at early time points, followed by decreased SOX2 expression and reduced cell viability at later time points in the corresponding cell types (**Fig. 2E**). Importantly, cell death was strictly dependent on the presence of the targeted interaction (**Fig. 2E**, WST-1), confirming its functional necessity.

We next examined the cell-type specificity of ASO-mediated targeting of RNA interactions using a multicellular human organoid glioma model. This platform was established by co-culturing patient-derived GSCs with iPSC-derived cortical brain organoids that recapitulate major neural lineages, including mature neurons, astrocytes, oligodendrocytes, and intermediate progenitors such as oligodendrocyte precursor cells and early neural precursors^17^ **(Supplemental Figure 3)**. To mimic the native tumor microenvironment, we incorporated myeloid cell component at the onset of organoid formation by combining iPSCs engineered for a myeloid fate with uncommitted iPSCs. This approach allowed the myeloid cells to progressively differentiate, migrate, and integrate into the maturing tissue, resulting in engrafted microglia-like cells that recapitulate endogenous cell-cell interactions and the complex architecture of the cortical organoid. Using ASOs carrying UV-cleavable molecular barcodes (**Fig. 2F**), we first quantified ASO uptake across organoid cell populations (**Fig. 2G, top**). Although the ASO targeting the glioma-specific LINC01206–SOX2 interaction was broadly taken up by all cells within the organoids, it selectively eliminated glioma cells while sparing non-malignant populations (**Fig. 2G, bottom;** scRNA-Seq).

Similarly, we identified glioma-specific lncRNA interactors for additional essential oncogenes, including EGFR, TERT, and CDK6 (**Supplemental Fig. 2**). ASO-mediated targeting of these RNA interactions likewise disrupted RNA–RNA and DNA–DNA contacts at early time points, followed by suppression of the corresponding oncogenes and glioma-specific cell death at later time points.

Together, these findings establish cell-type-specific RNA interactions as a therapeutically actionable and highly selective class of targets.

## SCIENCE-seq identifies glioma-specific cis-regulatory DNA elements controlling oncogene expression

We next asked whether DNA SCIENCE could resolve functional cis-regulatory interactions governing oncogene expression in a cell-type-specific manner. We focused on CD44, a well-established glioblastoma driver that promotes invasion, stem-like cell maintenance, and therapeutic resistance^18,19^.

Bulk RNA-seq analysis revealed that CD44 expression is minimal in myeloid cells, neurons, and oligodendrocytes, moderate in astrocytes and endothelial cells, and markedly elevated in OPCs and GSCs (**Fig. 3A**). Inspection of the CD44 locus identified multiple flanking cis-regulatory elements marked by H3K27ac, suggestive of active enhancers (**Fig. 3B**).

**Figure 3.**
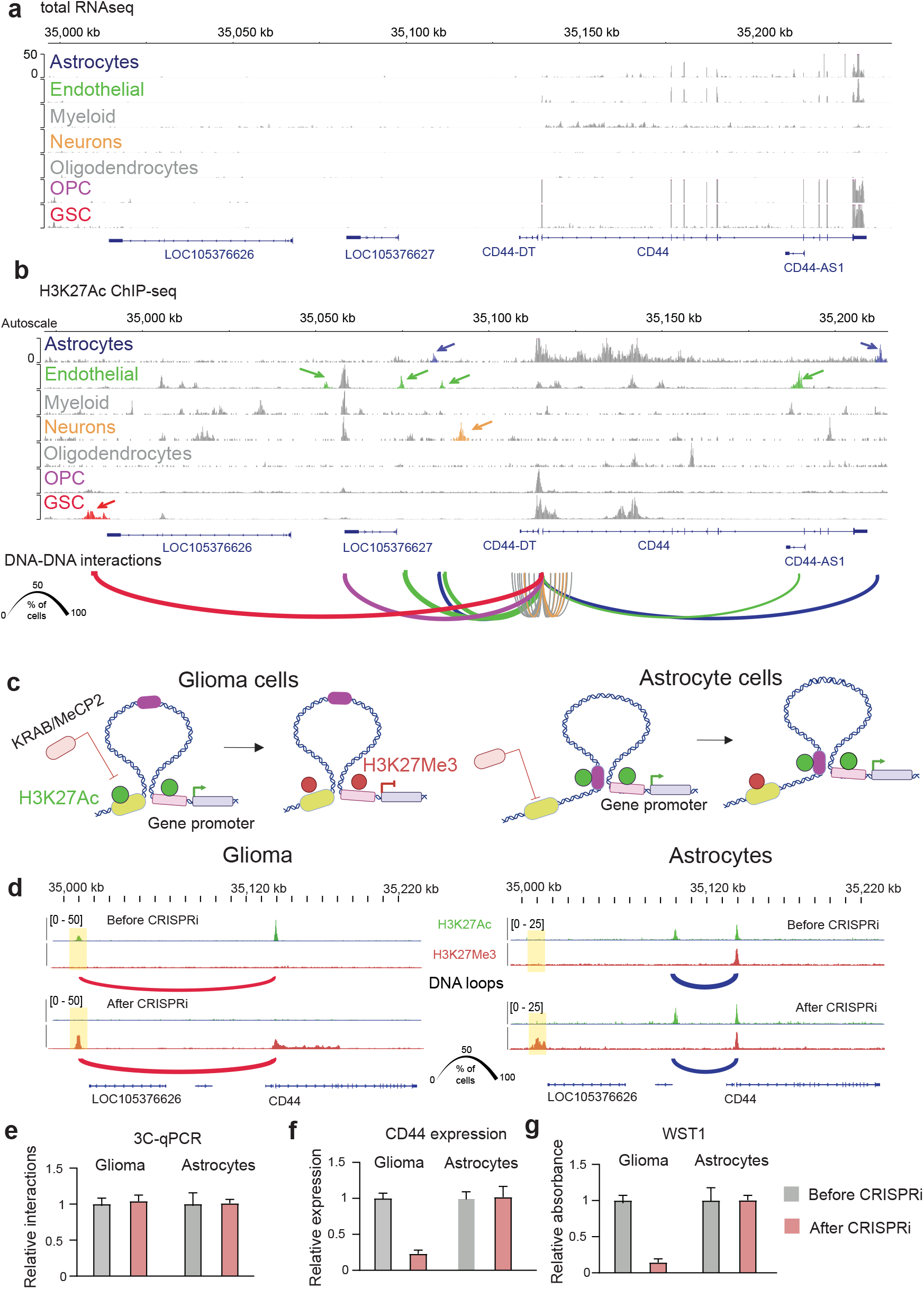
DNA SCIENCE reveals glioma-specific cis-regulatory elements. a. Total RNA-seq coverage of the CD44 locus across six distinct normal brain cells and glioma stem cells. b. H3K27Ac ChIP-Seq peaks marking active cis-regulatory elements surrounding CD44 across six distinct normal brain cells and glioma stem cells. Cell-type specific elements are highlighted by color and arrow. DNA–DNA interactions between CD44 promoter and those elements are shown as arcs, color-coded by cell type. Arc width represents the percentage of cells with a corresponding DNA-DNA interaction. c. Schematic of the CRISPRi (dCas9-KRAB-MeCP2) strategy for targeted epigenetic silencing of a glioma-specific CD44 enhancer. While both glioma and astrocyte lineages express CD44, its promoter is physically tethered to distinct cell-type-specific enhancers. CRISPRi-mediated silencing of the glioma-specific enhancer propagates to the CD44 promoter via pre-established DNA loops in glioma, whereas the CD44 locus in astrocytes remains unaffected due to its alternative DNA loop. d. ChIP-seq tracks illustrate an “epigenetic switch” at the CRISPRi-targeted glioma-specific enhancer, characterized by the loss of H3K27ac (active) in glioma and gain of H3K27me3 (repressed) in both glioma (left) and astrocytes (right). Arc width represents the percentage of cells with a corresponding DNA-DNA interaction. e. Chromatin DNA loops (arcs, taken from Fig. 3D) remain stable in both glioma and astrocytes following CRISPRi-mediated silencing. Data represent 3C-qPCR measurements of corresponding interaction frequencies (n = 3; mean with SD). f. CD44 mRNA expression measured by RT-qPCR in GSC (n=3, mean with SD) or Astrocytes (n=3; mean with SD) following CRISPRi. g. Cell death measured by WST1 assay in Glioma stem cells (n=3; mean with SD) and astrocytes following CRISPRi (n=3; mean with SD).

DNA SCIENCE demonstrated that the CD44 promoter engages in highly specific, cell-type-dependent chromatin interactions with these elements (**Fig. 3B, bottom arcs**). Notably, in GSCs- but not in other cell types- the CD44 promoter interacts with a distal regulatory element located approximately 150 kb upstream and enriched for H3K27ac, supporting its function as a glioma-specific enhancer.

To functionally validate this interaction, we employed CRISPR interference (dCas9-KRAB-MeCP2) to silence this candidate enhancer by inducing a repressive chromatin state (**Fig. 3C**). Targeted silencing of this regulatory element induced the deposition of H3K27me3 in both glioma and astrocytes **(Figure 3C**, highlighted in yellow); however, the gain of H3K27me3 at the *CD44* promoter occurred exclusively in glioma. Notably, 3C-qPCR analysis revealed that cell-type-specific chromatin loops (**Figure 3D**, colored arcs) remained stable following CRISPRi in both lineages (**Figure 3E**). Functionally, CRISPRi-mediated silencing resulted in significant *CD44* downregulation and subsequently reduced glioma cell viability, whereas astrocytes remained viable (**Fig. 3F, G**). These data suggest that stable chromatin loops may serve as structural scaffolds that facilitate the exchange of epigenetic regulators between distal elements and their target promoters.

In summary, DNA SCIENCE can identify functionally relevant, cell-type-specific cis-regulatory interactions that control oncogene expression and represent potential therapeutic targets.

## SCIENCE-seq links RNA–RNA interactions to alternative splicing and transcript architecture

We next investigated whether RNA–RNA interactions contribute to the regulation of mRNA isoform diversity in glioma. To this end, we integrated SCIENCE-seq data with multiple RNA-seq datasets from glioma and normal brain tissues, including both bulk samples and purified cell populations.

Using the Whippet pipeline^20^, we identified widespread alterations in alternative splicing (AS) and transcript processing (TP) in glioma and GSCs relative to normal brain cells. The most prominent changes involved alternative terminal exon usage (TE) with alternative polyadenylation (APA), and alternative transcription start sites (TS), consistent with known oncogenic transcript remodeling^21^ (**Fig. 4A, B; Supplemental Figure 4 for additional examples**). To directly test the role of RNA–RNA interactions in splicing regulation, we focused on CDC42, a Rho GTPase implicated in glioma proliferation and survival^22^. TCGA analysis revealed two major CDC42 isoforms: a longer isoform (ENST00000344548) and a shorter isoform (ENST00000315554), differing in exon 3 inclusion and terminal exon usage (**Fig. 4C**). The longer isoform is strongly enriched in glioma (**Fig. 4C**) and has been shown to be more stable, thereby contributing to elevated CDC42 expression in GBM^23^.

**Figure 4.**
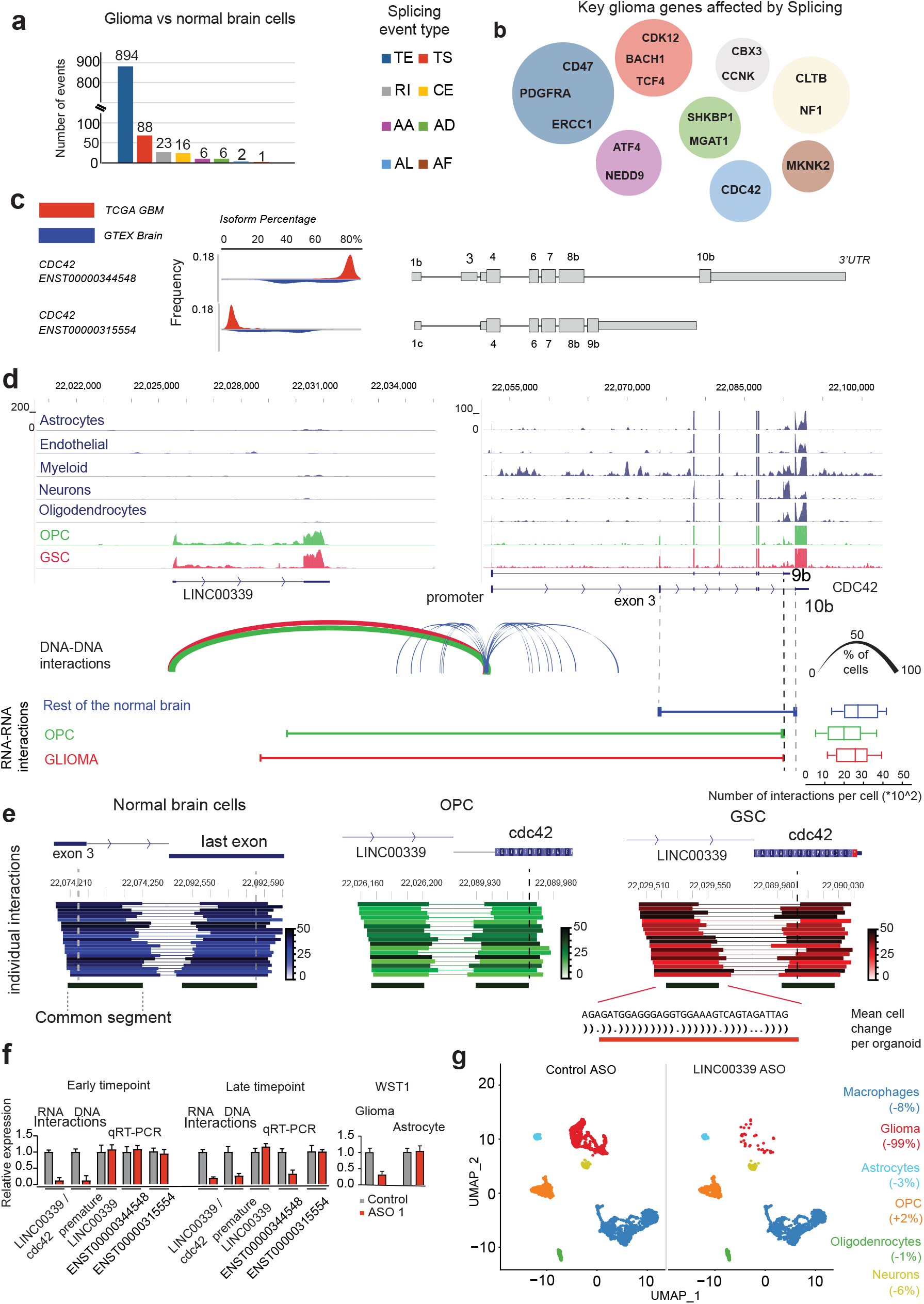
Combined RNA and DNA interactomics correlate with cancer-specific alternative splicing in glioma. a. Differential splicing landscape in glioma. Analysis of alternative splicing events between glioma and normal brain cell types using bona fide datasets. Bar graphs represent the frequency of specific splicing events (e.g., exon skipping), color-coded by event type. b. Regulation of glioma-specific splicing via RNA and DNA interactions. Statistically significant glioma oncogenes grouped by splicing event (as in Figure 4A). c. Isoform switching of CDC42 in glioma. Comparative analysis of CDC42 isoform percentage in glioma samples (TCGA, n=170) versus normal brain tissue (GTEx, n=870). Schematic representations of the corresponding transcript models are provided on the right. d. Splicing regulation of the CDC42 locus. Total Bulk RNA-Seq in different cell types shows cell type-specific lncRNA expression (top) and DNA or RNA common interacting segments measured by SCIENCE-Seq (bottom). Arc width (bottom, right) represents the percentage of cells with a corresponding DNA-DNA interaction. Boxplots (bottom, right) quantify the frequency of each RNA–RNA interaction per cell. e. Mapping of individual interacting RNA segments for therapeutic targeting. Intragenic localization and frequencies of individual interacting RNA segments across OPCs, Astrocytes, Oligodendrocytes and Glioma. The visualization highlights a glioma-specific common segment and aligned sequence (red) utilized for the design of Antisense Oligonucleotide (ASO) therapeutics. f. Targeted disruption of the glioma RNA and DNA interactome with ASOs. Treatment of glioma stem cells with ASO disrupts glioma-specific RNA (SPLASH-qPCR; n=3, mean with SD) and DNA (3C-qPCR; n=3, mean with SD) interactions between CDC42 and LINC00339, leading to decreased ENST00000344548 isoform expression (qRT-PCR; n=3, mean with SD) and glioma cell death (WST1; n=3, mean with SD). Measurements were taken in 2 timepoints g. LINC00339 ASO treatment induces selective glioma cell death. Organoids (n=2) were treated with Control ASO or LINC00339 ASO. Synchronized and overlayed UMAP plots display cell clusters following treatment with Control or LINC00339 ASO. Mean cell count changes per organoid (Right). Every dot is an individual cell.

We observed that the lncRNA LINC00339 is expressed specifically in GSCs and OPCs, but not in other brain cell types **(Fig. 4D)**. DNA SCIENCE **(Fig.4D, arcs)** revealed promoter–promoter interactions between LINC00339 and CDC42 selectively in these cell populations. Strikingly, RNA SCIENCE **(Fig 4D, connected bars)** identified direct interactions between premature LINC00339 and CDC42 RNA at exon 9b, an exon that is preferentially skipped in glioma cells.

In contrast, in normal brain cells lacking LINC00339 expression, CDC42 transcripts exhibited only intra-molecular RNA interactions, particularly between alternative exon 3 and downstream exon 10b, consistent with a transcript conformation favoring exclusion of both exons. These observations suggest that LINC00339 binding to CDC42 pre-mRNA promotes selection of specific terminal exons and drives glioma-specific splicing patterns.

To test this mechanism, we identified a glioma specific, high-occupancy CDC42-interacting region within LINC00339 (**Fig. 4E, red**) and designed a steric-blocking ASO targeting this site (**Fig. 4E, red line**). Disruption of this interaction in GSCs resulted in a rapid loss of RNA-RNA and DNA-DNA interactions between LINC00339 and CDC42, followed by a reduction in the tumor-associated long CDC42 isoform and reduced glioma viability at later time points **(Fig. 4F)**, supporting a causal role for RNA–RNA interactions in splicing regulation. Single-cell RNA sequencing (scRNA-seq) in human organoids demonstrated that the ASO targeting this glioma-specific interaction exhibits selective glioma toxicity (**Fig. 4G**).

## RNA–RNA interactions regulate alternative polyadenylation in glioma

Alternative polyadenylation (APA) is a widespread feature of cancer transcriptomes, often resulting in 3′UTR shortening and enhanced oncogene activity^24,25^. Consistent with this, our analysis identified TE/APA as the most significant transcript processing alteration distinguishing glioma from normal brain (**Fig. 4A**, **5A**).

**Figure 5.**
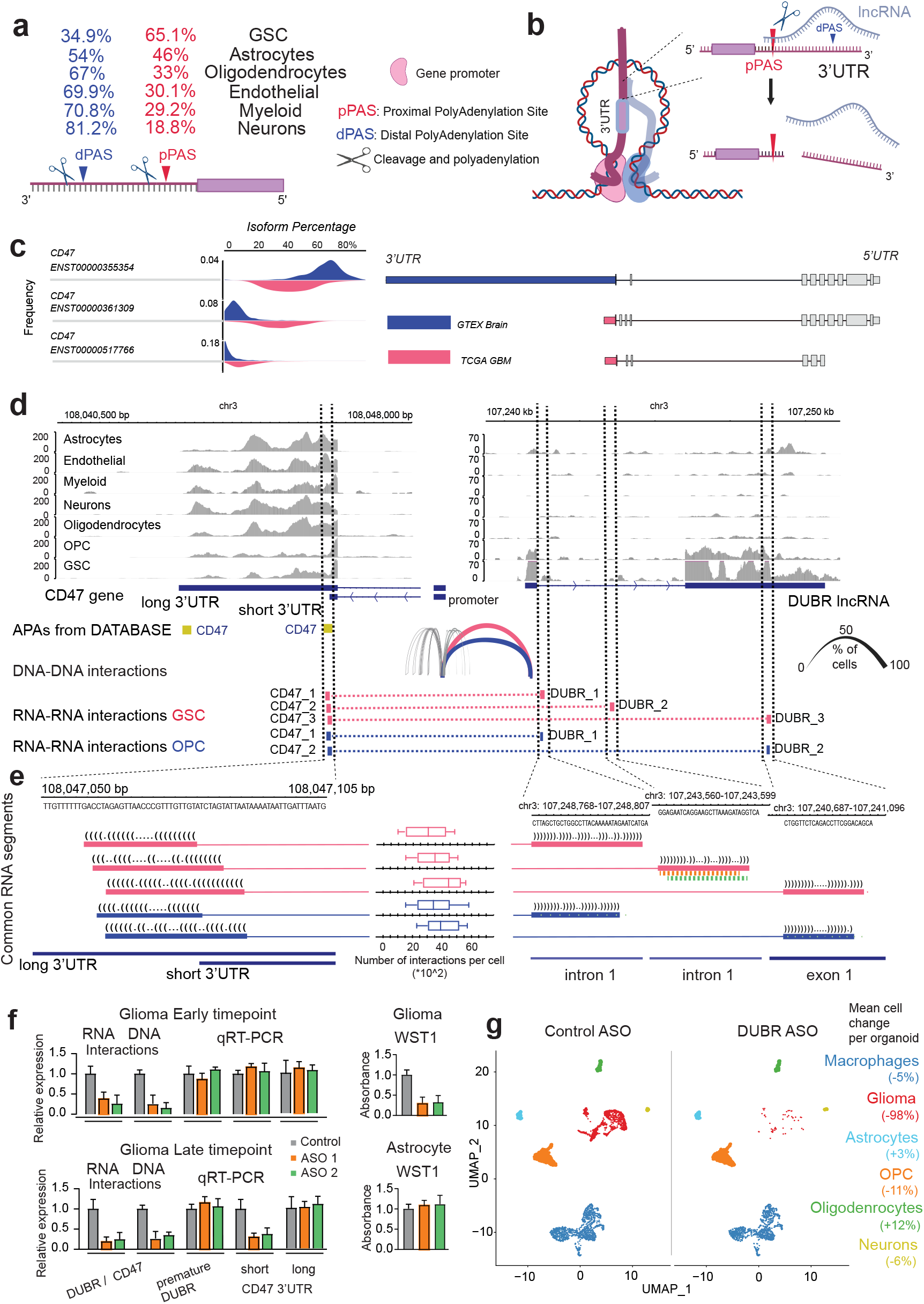
RNA and DNA interactomics define the choice for cell-type specific Alternative Polyadenylation. a. Global 3′UTR shortening in glioma. Comparative analysis of 3′UTR length across glioma and healthy brain cell types. The percentage of genes with short (pPAS) and long (dPAS) 3’UTRs are visualized. b. Mechanistic model of lncRNA-directed 3′UTR shortening. Synergistic DNA-DNA promoter contacts and nascent RNA-RNA interactions at the proximal polyadenylation site (pPAS) drive precise transcript cleavage resulting in 3′UTR shortening in glioma. c. Differential Alternative Polyadenylation of CD47 in malignant versus normal tissue. Comparative analysis of the percentage of short (ENST00000361309 and ENST00000517766) and long cd47 (ENST00000355354) isoforms were measured using TCGA glioma samples (n=170) and normal brain tissue (n=870). Schematic representations of the corresponding transcript models are provided on the right. d. Mechanistic basis for CD47 3′UTR truncation. Total RNA-seq read coverage across the 3’ UTR reveals a sharp decline at the proximal polyadenylation site (pPAS) specifically in GSCs and OPCs (left), which correlates with elevated DUBR lncRNA expression in these cells (right). Genomic annotations of alternative polyadenylation (APA) sites and cell type-specific DNA–DNA (DNA SCIENCE; arcs, which width (bottom, right) represents the percentage of cells with a corresponding DNA-DNA interaction) and RNA–RNA interactions (RNA SCIENCE; dotted lines) for GSCs and OPCs are provided below. e. Mapping of interacting RNA segments for therapeutic targeting. Common cell-type specific RNA segments for pPAS of CD47 and DUBR were visualized with their sequences, genomic positions and interaction counts per cell (RNA SCIENCE; boxplots, middle). Two ASO candidates targeting a glioma-specific DUBR intronic segment are indicated by colored dotted lines. f. Targeted disruption of the glioma RNA-RNA interactome with ASOs. Treatment of glioma stem cells with 2 ASO disrupts glioma-specific RNA-RNA (SPLASH-qPCR; n=3, mean with SD) and DNA-DNA (3C-qPCR; n=3, mean with SD) interactions between CD47 and DUBR at early timepoint, leading to decrease of shorter (oncogenic) isoform expression (qRT-PCR; n=3, mean with SD) and cell death (WST1; n=3, mean with SD) at late timepoint. Astrocyte cells stay intact. g. DUBR ASO treatment induces selective glioma cell death. Organoids (n=2) were treated with Control ASO or DUBR ASO (ASO1 from Figure 5F). Synchronized UMAP plots display cell clusters following treatment with Control or DUBR ASO. Mean cell count changes per organoid (Right). Every dot is an individual cell.

RNA SCIENCE data suggest that APA site selection is mediated by interactions between pre-mRNAs and intronic lncRNAs precisely at Proximal Polyadenylation Site (**Fig. 5B, Supplemental Figure 5 shows additional examples**). A representative example is CD47, which expresses multiple isoforms with distinct 3′UTRs that influence protein localization and function^26^.

Analysis of TCGA datasets revealed that the long CD47 isoform (ENST00000355354) predominates in normal brain, whereas shorter isoforms (ENST00000361309 and ENST00000517766) are enriched in glioma. Bulk RNA-seq coverage confirmed preferential usage of a proximal APA site in OPCs and GSCs (**Fig. 5D**).

DNA SCIENCE revealed strong promoter–promoter interactions between CD47 and the lncRNA DUBR specifically in OPCs and GSCs **(Figure 5D, arcs)**, whereas in normal brain cells CD47 exhibited weak and diffuse chromatin contacts with the neighboring areas. RNA SCIENCE further identified direct RNA-RNA interactions between nascent DUBR and the proximal APA region of CD47 transcripts in glioma cells (**Fig. 5E**).

Targeting the GSC-specific interaction with ASOs directed against the DUBR interacting region disrupted RNA–RNA and DNA-DNA contacts and shifted CD47 isoform usage toward the longer 3′UTR, resulting in selective glioma cell death (**Fig. 5F**). Consistently, scRNA-seq analysis in human organoids demonstrated that ASO-mediated targeting of this glioma-specific interaction induces selective glioma toxicity while sparing non-malignant cells (**Fig. 5G**). Together, these findings demonstrate that RNA–RNA interactions can directly govern APA site selection and constitute a previously unrecognized mechanism of oncogenic transcript remodeling.

## HOX lncRNAs establish a glioma-specific trans-chromosomal regulatory hub

A striking and unexpected finding from SCIENCE-seq was the identification of a higher-order, inter-chromosomal interactome spanning all four HOX clusters. HOXA, HOXB, HOXC, and HOXD clusters with embedded lncRNAs are activated in GBM (TCGA, n = 170), contrasting with their near-complete repression in normal brain tissue (TCGA, n = 870) (**Fig. 6A**). Across tumors, expression of HOX coding and non-coding transcripts was highly correlated, suggesting the existence of a shared regulatory architecture (**Fig. 6B**).

**Figure 6.**
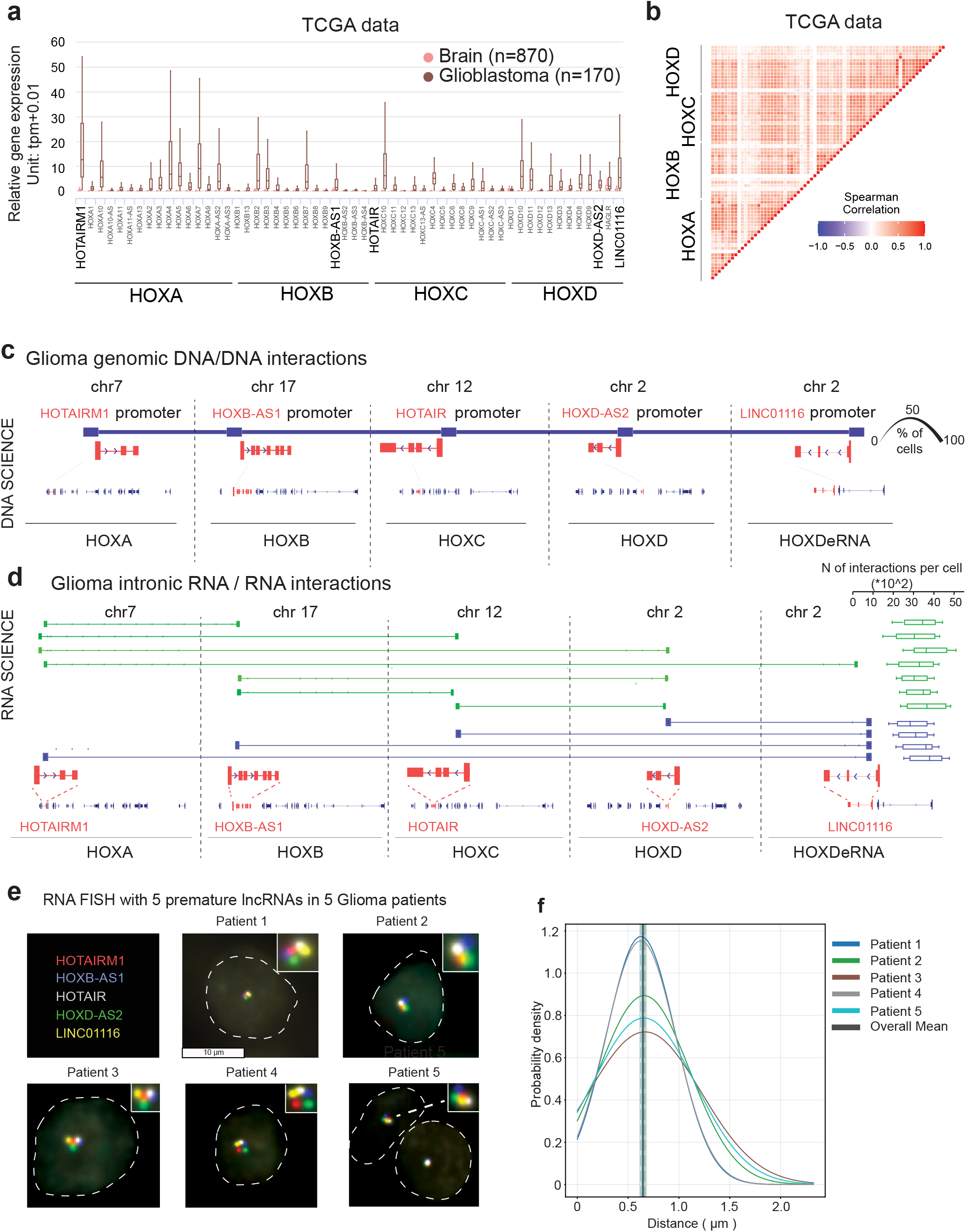
HOX gene clusters form an inter-chromosomal transcriptional hub in glioma. a. Overexpression of the HOX transcriptional program in glioma. Box plots illustrate the significant enrichment of HOX family mRNA levels in Glioblastoma (TCGA, GBM; n=170) compared to healthy brain tissue (GTEx, n=870). Highlighted in UPPERCASE are specific lncRNAs embedded within the HOX genomic clusters. b. HOX genes correlate in their expression. Heatmap of Pearson correlation coefficients for pairwise HOX mRNA expression levels (TCGA, GBM; n=170). c. HOX lncRNA co-interacting with their promoters. DNA-DNA interaction analysis (DNA SCIENCE) identifies 5 lncRNA co-interacting with their promoters in a single read at a single cell resolution. Line width represents the percentage of cells with a corresponding DNA-DNA interaction (bottom right). d. HOX premature lncRNAs form an inter-chromosomal hub in glioma. Pairs of interacting common intronic segments (RNA SCIENCE; bars connected with lines) from HOX premature lncRNAs (green) are aligned to their respective transcript models (red, bottom). HOXDeRNA interactions are highlighted in blue. The abundance of each pair of interactions was quantified at the single-cell level in glioma (boxplot, right). e. Validation of the HOX lncRNA hub in primary tumors. Sequential single-molecule Fluorescence In Situ Hybridization (smFISH) in primary glioma tissue (n=3) confirms the co-localization of five premature lncRNAs originating from four HOX clusters. f. Spatial clustering of 5 HOX premature lncRNAs in primary glioma samples. Histogram displaying the distribution of physical distances between pairs of premature HOX lncRNAs across five glioma patients.

DNA SCIENCE analysis of primary glioma samples demonstrated that these four HOX loci - located on distinct chromosomes-physically co-interact in glioma cells. The interacting genomic regions mapped specifically to the promoters of five lncRNA genes embedded within HOX clusters: **HOTAIRM1 (HOXA), HOXB-AS1 (HOXB), HOTAIR (HOXC), HOXD-AS2 (HOXD), and LINC01116 (**or HOXDeRNA, **HOXD** enhancer) (**Fig. 6C**). Notably, 95% of glioma cells harbor at least one concatenate DNA read containing all five lncRNA promoters, indicating that these regions co-exist within shared nuclear complexes analyzed at single-cell resolution. In contrast, such inter-chromosomal contacts were not detected in normal brain cells.

In parallel, RNA SCIENCE identified direct RNA–RNA interactions among the nascent transcripts of these five lncRNAs, with interacting segments mapped to discrete regions within each transcript, as detailed in **Fig. 6D**. Importantly, we did not detect comparable interactions between protein-coding HOX pre-mRNAs across clusters, indicating that this architecture is specifically mediated by lncRNAs rather than coding transcripts. Together, these data define a glioma-specific, lncRNA-centered, trans-chromosomal structure that we term a **HOX interactomic hub**.

To independently validate this architecture, we performed smFISH using intronic probes targeting the five HOX lncRNAs in patient-derived GBM tissues (**Fig. 6E**). We detected 1–2 foci per lncRNA per cell, demonstrating that these intronic smFISH probes predominantly capture nascent transcripts accumulating at their transcription sites, consistent with previous reports^27,28^. Quantification of inter-probe distances revealed consistent spatial co-localization of these nascent transcripts across cells and patients, with minimal variability between lncRNA pairs (**Fig. 6F**), supporting the existence of a stable, higher-order nuclear assembly.

## Nascent lncRNAs nucleate and maintain the HOX trans-chromosomal hub

We next sought to determine whether the HOX interactomic hub is driven by protein scaffolding or by RNA itself. To this end, we performed bulk 4C-seq using each of the five HOX lncRNA promoters as viewpoints. These experiments confirmed reciprocal inter-chromosomal interactions among all five loci in glioma cells.

Strikingly, treatment with Proteinase K did not disrupt these interactions, whereas pooled siRNA-mediated depletion of the five HOX premature lncRNAs abolished them (**Fig. 7A**), demonstrating that the integrity of the HOX hub depends on nascent RNA transcripts rather than protein-mediated chromatin organization.

**Figure 7.**
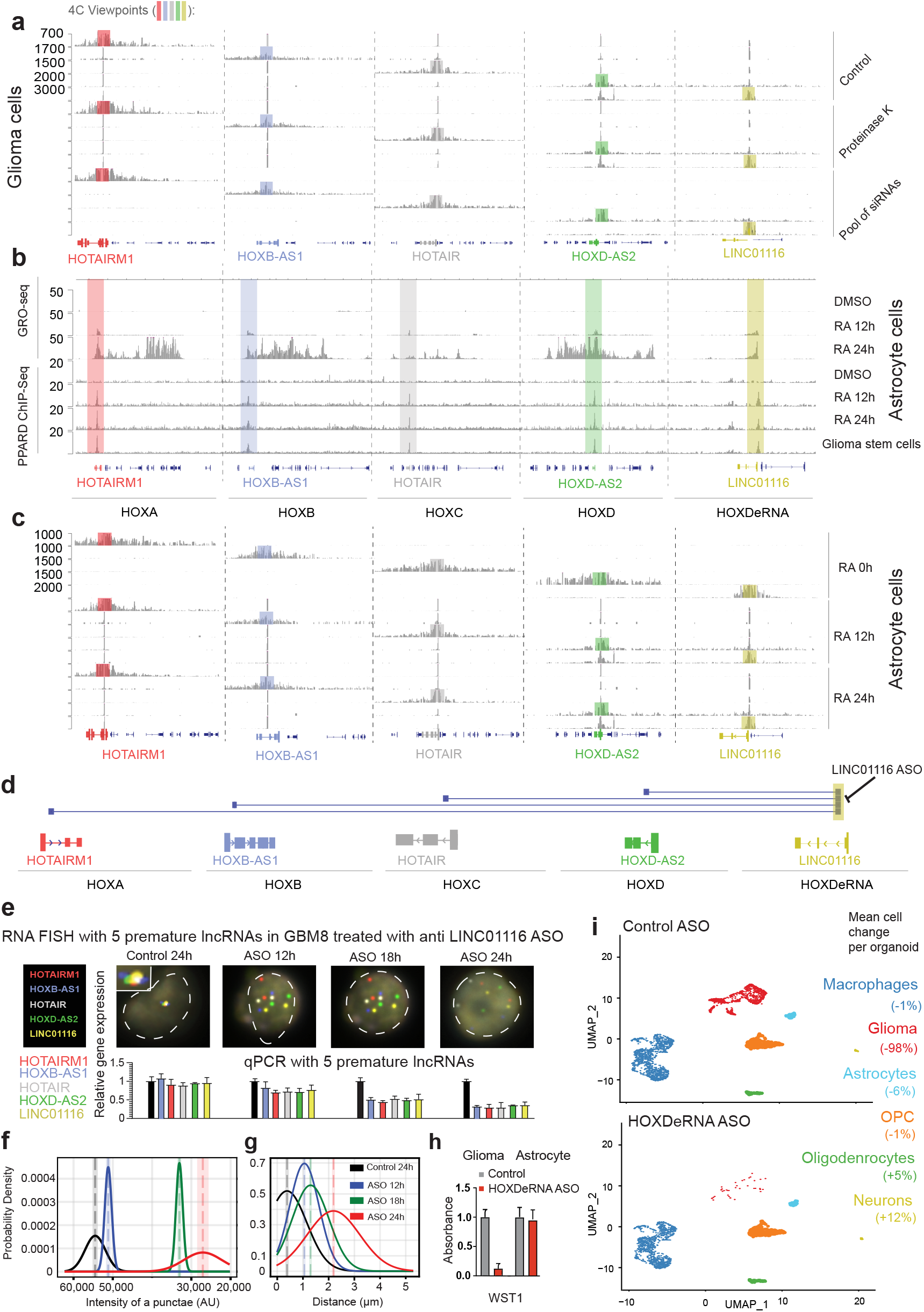
Regulation and therapeutic disassembly of the HOX transcriptional hub. a. Chromatin DNA-DNA interactions between all 4 HOX clusters strongly depend on premature lncRNAs, but not on proteins. Bulk 4C-seq analysis in GSCs (GBM8, n=2) with viewpoints at 5 HOX lncRNAs (highlighted with color) performed in Control, Proteinase K and 5 premature HOX lncRNA combined siRNA KD. b. De novo assembly of the HOX hub following ligand-induced activation. Bulk 4C-Seq with 5 viewpoints (highlighted with color) taken at 5 HOX lncRNAs shows the formation of reciprocal inter-chromosomal DNA-DNA contacts during ATRA treatment of astrocytes. These interactions are absent in control (DMSO) condition but are established within 12h of treatment. c. Temporal dynamics of ATRA-induced HOX activation in human astrocytes. Treatment with All-Trans Retinoic Acid (ATRA) triggers sequential transcriptional activation of the HOX clusters, with embedded lncRNAs preceding protein-coding gene expression (Gro-Seq, HOX lncRNAs loci are highlighted).ChIP-seq profiles at 12h and 24h demonstrate that PPARD selectively binds HOX lncRNA loci early in the activation phase, mirroring the binding pattern observed in Glioma Stem Cells (PPARD ChIP-Seq). d. Diagram showing intronic RNA-RNA interactions between HOXDeRNA and other premature lncRNAs. A common segment within LINC01116/HOXDeRNA is highlighted in gold. An ASO was synthesized to selectively target this sequence. e. The disruption of the HOX transcriptional hub with ASO. Sequential smFISH imaging and RT-qPCR (n=3, mean with SD) of 5 nascent HOX lncRNAs in Glioma Stem Cells (GSCs) over a 24-hour time course. f. Intensities of premature HOX lncRNAs gradually decline after HOX hub disruption. Distribution of HOX lncRNAs intensities were measured for Fig. 7E and visualized as Gaussian probability densities for each time-point (colored lines). g. Distances between premature HOX lncRNAs gradually increase after HOX hub disruption. Pairwise distances between HOX premature lncRNA transcripts were measured for Figure 7E and visualized as Gaussian probability densities for each time-point (colored lines). h. ASO treatment induces selective cell death in glioma cells, but not in astrocytes. WST1 assay was performed in glioma (n=3, mean with SD) and astrocytes (n=3, mean with SD). i. HOXDeRNA ASO treatment induces selective glioma cell death. Organoids (n=2) were treated with Control ASO or HOXDeRNA ASO. Synchronized UMAP plots display cell clusters following treatment with Control or HOXDeRNA ASO. Mean cell count changes per organoid (Right). Every dot is an individual cell.

## HOX hub formation is inducible and precedes cluster-wide transcriptional activation

To determine whether HOX hub formation is a cause or consequence of transcriptional activation, we utilized an inducible system based on all-trans retinoic acid (RA), a known activator of HOX gene expression^29^. Immortalized human astrocytes (NHAs), which lack baseline HOX expression, were treated with RA and analyzed over time.

GRO-seq revealed that transcription is first initiated at discrete genomic intervals corresponding to the HOX lncRNA promoters-identified here as **“Hot Spots”**-within 12 hours of RA treatment, followed by activation of entire HOX clusters by 24 hours (**Fig. 7B, top**). Thus, transcriptional activation originates at the same loci that anchor the HOX hub.

ChIP-seq analysis of PPARD, a nuclear receptor and transcription factor functionally activated by FABP5-mediated retinoic acid (RA) delivery in oncogenic contexts^30^, revealed strong enrichment at these Hot Spots following RA treatment, closely mirroring the binding profile observed in glioma stem cells (**Fig. 7B, bottom**). These findings suggest that oncogenic signaling converges on these loci as an initiating mechanism for hub formation.

Using 4C-seq, we found that prior to RA induction, HOX lncRNA loci interact primarily within their local chromosomal neighborhoods. However, as early as 12 hours after RA treatment-coincident with initial lncRNA transcriptional activation, and prior to activation of the entire HOX regions-reciprocal inter-chromosomal contacts among HOX loci emerge (**Fig. 7C**).

These results indicate that **formation of the trans-chromosomal HOX hub is an early and regulated event that precedes full transcriptional activation**, rather than a passive consequence of gene expression.

## Targeted disruption of HOX lncRNA interactions dismantles the hub and suppresses transcription

To determine whether the HOX hub is functionally required for transcription, we designed a steric-blocking ASO targeting the LINC01116/HOXDeRNA intronic interaction interface identified by RNA SCIENCE (**Fig. 7D, highlighted in gold**). This region serves as a shared contact site mediating interactions between LINC01116 and four additional HOX lncRNAs.

Glioma cells were treated with the ASO, and hub dynamics were monitored by sequential smFISH imaging and RT-qPCR over a 12-, 18-, and 24-hour time course. ASO treatment induced rapid hub disassembly, accompanied by a reduction in nascent lncRNA transcripts (**Fig. 7E, F**). This effect was highly coordinated across the network, as evidenced by concordant decreases in both smFISH fluorescence intensity and RT-qPCR measurements for all five HOX lncRNAs. Consistent with progressive hub disruption, the physical distances between HOX lncRNA transcription sites increased over time, indicating a loss of higher-order spatial organization (**Fig. 7G**).

To assess the functional consequences of hub disruption, we examined cell viability following ASO treatment. WST-1 assays demonstrated potent glioma cell killing with no detectable toxicity to healthy astrocytes (**Fig. 7H**). Furthermore, scRNA-seq analysis in human organoid models showed that targeting this glioma-specific RNA interaction selectively eliminated glioma cells while sparing non-malignant cell populations (**Fig. 7I**).

Together, these findings demonstrate that a discrete RNA interaction interface is essential for maintaining a higher-order trans-chromosomal regulatory hub that coordinates transcription across multiple genomic loci. Disruption of this interface collapses the hub, synchronously silences its constituent lncRNAs, and selectively impairs glioma cell viability.

## METHODS

### Cells and Cell Lines

All studies involving human cells were conducted in accordance with the ethical guidelines and policies of the Institutional Review Board (IRB) at **Brigham and Women’s Hospital**.

#### Glioma Stem-like Cells (GSCs)

Low-passage, patient-derived glioblastoma stem-like GBM8 cells were kindly provided by Dr. Hiroaki Wakimoto (Massachusetts General Hospital). The genetic, molecular, and tumorigenic profiles of the GBM8 line have been previously characterized and validated^31,32^. GBM8 were maintained as neurospheres in serum-free Neurobasal medium (Gibco) supplemented with: B-27 Plus and N-2 supplements (Gibco), 3 mM GlutaMAX (Gibco), 20 ng/mL recombinant human FGF2 and EGF (Sigma-Aldrich), 2 μg/mL heparin (Sigma-Aldrich), 50 units/mL penicillin/streptomycin (Gibco). Cells were passaged upon reaching a diameter of approximately 200–300 μm via chemical dissociation using the Neurocult Stem Cells Dissociation Kit (Stem Cell Technologies) to ensure high viability and maintenance of the undifferentiated state.

#### Immortalized Normal Human Astrocytes

(NHA): NHA, immortalized via E6/E7 and hTERT expression (a generous gift from Dr. Yukihiko Sonoda), served as non-malignant controls. These cells were cultured in **Astrocyte Medium** (ScienCell, # 1801) according to the manufacturer’s instructions.

Cell cultures were periodically tested for mycoplasma

### Organoid generation

Glioma organoids were generated with modifications to previously described protocols^17^. To incorporate myeloid cells into the organoid architecture, human induced pluripotent stem cells (iPSCs) were engineered to express PU.1 under a tetracycline-inducible (Tet-On) promoter. Briefly, wild-type (WT) iPSCs were transfected using a Lonza 4D-Nucleofector System with the P3 Primary Cell 4D-Nucleofector X Kit (Lonza) by co-transducing a piggyBac transposon plasmid containing PU.1 (Addgene, #179514) and a plasmid encoding the piggyBac transposase at a 3:1 ratio (1.5 µg transposon: 0.5 µg transposase). For organoid generation, WT iPSCs were differentiated into the neuroectoderm lineage to form the neural component, while the engineered PU.1-iPSCs were differentiated into the mesoderm lineage to form the myeloid component. Both cell lines were cultured in parallel for 3 days in a basal differentiation medium consisting of DMEM/F12 supplemented with 2% B27, 1% N2, 1% non-essential amino acids, and 0.2% penicillin/streptomycin. For neuroectoderm commitment, the basal medium was supplemented with 10 µM SB431542 (MedChemExpress #HY-10431), 0.25 µM LDN-193189 (MedChemExpress #HY-12071), and 0.1 µM retinoic acid (Sigma-Aldrich). Concurrently, mesoderm differentiation was induced by supplementing the basal medium with 100 *ng*/*mL* BMP4 (BioLegend, #595202), 100 *ng*/*mL* VEGFA-165 (BioLegend, #583702), and 20 *ng*/*mL* FGF2 (BioLegend #710304).

Following the 3-day lineage-specific induction, neuroectoderm and mesoderm colonies were detached using Dispase II (Sigma-Aldrich #SCM133) and mechanically dissociated via gentle pipetting. To assemble the co-culture spheroids, the dissociated colonies were combined at a 9:1 ratio (9 parts neuroectoderm to 1 part mesoderm) and seeded into ultra-low attachment 96-well plates (Corning). Organoids were maintained as previously described, with the culture medium supplemented with 1 µM doxycycline hydrochloride (Sigma-Aldrich) for the initial 30 days, followed by 3µM doxycycline hydrochloride for an additional 30 days to drive PU.1 expression over a total induction period of 60 days. Following the 60-day induction phase, doxycycline was withdrawn from the culture medium, and the organoids were allowed to mature for an additional 15 days prior to glioma stem cell (GSC) engraftment in Neurobasal medium supplemented with 2% B27, 1% N2, 1% NEAA, and 0.2% P/S.

For GSC engraftment, 10 matured organoids were transferred into a single well of an ultra-low attachment 6-well plate containing 2 mL of organoid medium. Subsequently, 100,000 GSCs pre-transduced with lentivirus expressing CFP were added to the well as a single-cell suspension. The organoids and GSCs were co-incubated for 24 h, after which the organoids were washed thoroughly to remove non-engrafted cells. The engrafted GSCs were then allowed to invade and integrate into the organoid architecture for an additional 21 days before experimental use.

### Organoid immunostaining

GSC-engrafted organoids were fixed in 4% paraformaldehyde for 1 h, cryoprotected in 30% sucrose (Sigma-Aldrich, #S0389-500G) until settled, embedded in OCT compound (Fisher, #23-730-571), and sectioned at 15 µm thickness. Slides were washed three times in TBS to remove OCT, permeabilized in TBS containing 0.5% Triton X-100 (Sigma-Aldrich #X100-500ML), and blocked for 1 h in TBS with 0.1% Triton X-100, 1% BSA (Sigma-Aldrich, #A2153-10G), and 5% normal goat serum (Vector Laboratories, #S-1000-20).

Primary antibodies were applied overnight at 4 °C against GFP (Antibodies Inc., #GFP-1010; used to detect CFP-expressing GSCs), βIII-Tubulin (CST, #5568S), GFAP (CST, #3670S), Olig1 (Sino Biological, #100577-T38), SOX2 (CST, #4900S), Olig2 (CST, #39588S), CNP (BioLegend, #836404), MBP (Abcam, #ab40390), and TMEM119 (BioLegend, #853302). Following washes, slides were incubated with corresponding secondary antibodies (goat anti-rabbit AF647 [#A-21245], goat anti-mouse AF555 [#A-11001], and goat anti-chicken AF488 [#A-11039]; ThermoFisher) together with Hoechst 34580 nuclear stain (1:1000; ThermoFisher, #H21486). Slides were mounted with ProLong Diamond (ThermoFisher, #P36970) prior to imaging on a Nikon W1-SoRa confocal microscope.

### GBM primary tumor samples

Clinical GBM specimens were obtained from Dana-Farber Cancer Institute, per a protocol approved by the Institutional Review Board. Detailed mutational spectrum—including nucleotide variants, copy number variants, and structural variants—are comprehensively summarized in **Supplemental Table 2**.

### siRNA transfection

Cells in 24 well plates were transfected with a pool of 5 siRNAs targeting 5 unspliced HOX-associated lncRNAs according to LipoRNAiMax protocol. siRNA sequences are listed in **Supplemental Table 3.**

### Cell viability assay (WST1)

Cell metabolism and viability were quantified by Cell Proliferation Reagent WST-1 (Cat# 5015944001, Sigma), according to manufacturer’s instructions.

### ASO design and transfection

Steric antisense oligonucleotides (ASOs) were designed as 20-nt single-stranded DNA oligonucleotides in which every nucleotide contained a 2′-O-methoxyethyl (2′-MOE) modification and all internucleotide linkages were phosphorothioate. ASO sequences are provided in **Supplemental Table 3**.

Barcoded ASOs (Supplemental Table 3) were synthesized with the following structure:

ACACTCTTTCCCTACACGACGCTCTTCCGATCTNNNNNNNNNNNNNNNNNNNNNNNNNGATCGGAAGAGCACACG/iPCLinker/[ASO]

where (N)\₍₂₅₎ denotes a 25-nucleotide random barcode, *iPCLinker* is an internal UV-cleavable linker, and *[ASO]* represents the 20-nt fully modified ASO sequence. The barcode region was positioned between TruSeq-compatible Read 1 and Read 2 adapter sequences to enable barcode identification by next-generation sequencing. The ASO portion consisted of 20 nucleotides carrying 2′-MOE modifications at every position and phosphorothioate linkages throughout.

### Lipid nanoparticle formulation for organoid transfection with ASO

Lipid nanoparticles (LNPs) were formulated using an ionizable lipid composition consisting of 50 mol% DLin-MC3-DMA (MC3; Cayman Chemical, Cat no# 34364), 38.5 mol% cholesterol (Sigma-Aldrich, Cat no# C8667), 10 mol% 1,2-distearoyl-sn-glycero-3-phosphocholine (DSPC; Avanti Polar Lipids, Cat no# 850365P-1g), and 1.5 mol% 1,2-dimyristoyl-rac-glycero-3-methoxypolyethylene glycol-2000 (DMG-PEG2000; Avanti Polar Lipids, Cat no# 880151P-1g). Lipid components were dissolved in absolute ethanol immediately prior to nanoparticle formulation.

Barcoded ASOs were dissolved in 50 mM sodium acetate buffer (pH 4.0; Thermo Fisher Scientific, Cat no# J60104.AK). The aqueous ASO solution was mixed with the ethanolic lipid solution at a volumetric ratio of 3:1 (aqueous), corresponding to an N/P ratio of 6, by vigorous vortex mixing to promote spontaneous nanoparticle self-assembly. The ASO concentration in the initial formulation mixture was approximately 0.05 mg/mL. Following mixing, the LNP formulation was incubated at room temperature for 15 min and then diluted with PBS to reduce the residual ethanol concentration to below 0.5% (v/v) prior to biological applications.

Human GBM organoids were treated with ASO-loaded LNPs at a final ASO concentration of 100 nM for 5 days, under standard culture conditions. Following treatment, the organoids were mechanically and chemically dissociated into single-cell suspensions using the NeuroCult™ Chemical Dissociation Kit for Mouse CNS Stem Cells (STEMCELL Technologies) according to the manufacturer’s instructions. Single-cell suspensions were subsequently processed for downstream single-cell analyses, including barcode detection and transcriptomic profiling.

### Bulk ChIP-Seq

Chromatin Immunoprecipitation (ChIP) assays were performed using the SimpleChIP® Enzymatic Chromatin IP Kit (Cell Signaling Technology, #9003) according to the manufacturer’s instructions. Briefly, 10 million cells were cross-linked with 1% formaldehyde for 10 minutes at room temperature, followed by quenching with glycine. Cells were washed twice with ice-cold PBS, and the resulting pellets were resuspended in RIPA buffer (Boston BioProducts, #BP-115X) supplemented with a protease inhibitor cocktail (Roche, 11836170001). Chromatin was fragmented to a median size of 300 bp using a MISONIX S-4000 Sonicator (30% amplitude, 30s ON/30s OFF cycles for 30 minutes). For each immunoprecipitation, 20 μg of sheared chromatin was diluted in 1 mL of IP Dilution Buffer (16.7 mM Tris-HCl pH 8, 167 mM NaCl, 1.2 mM EDTA, 1% Triton X-100, and 0.01% SDS) and incubated overnight at 4°C with 10 μg of anti-PPARD antibody (Abcam, #ab8937) or H3K27Ac (#4353, Cell Signaling) or H3K27Me3 (#9733, Cell Signaling). Immune complexes were captured by incubation with 30 μL of Dynabeads™ Protein G (Thermo Fisher) for 4 hours at 4°C. Beads were subjected to a stringent washing series: twice in low-salt buffer (150 mM NaCl), once in high-salt buffer (500 mM NaCl), and finally in TE buffer. Chromatin was eluted and de-crosslinked at 65°C overnight. Purified DNA was recovered using the Monarch PCR and DNA Cell Cleanup Kit (NEB, #T1030L). Library Preparation and Sequencing ChIP-seq libraries were constructed from purified DNA using the NEBNext® Ultra™ II DNA Library Prep Kit for Illumina® (NEB, #E7546S). Library quality and concentration were assessed via Agilent Bioanalyzer. Libraries were sequenced on the Illumina HiSeq 2500 platform to generate 50 – 150 bp single-end reads.

### ChIP-Seq Data Analysis

Low-quality reads (MAPQ<=30) were filtered out with BAMTools (v2.5.3). The remaining high-quality reads were aligned to the human reference genome (**GRCh38/hg38**) using **Bowtie2**^33^ (v2.5.3) with default end-to-end sensitive parameters. To mitigate potential amplification bias, PCR duplicates were computationally identified and removed from the aligned BAM files using **SAMtools**^34^ (*rmdup*, v1.13.7).

For visualization, aligned BAM files were converted to BigWig format using deepTools^35^ (v3.5.0) bamCoverage without additional scaling or normalization. All track data were visualized using the Integrative Genomics Viewer ^36^ (IGV, v2.16.0).

### CRISPR interference (CRISPRi)

The mammalian expression lentiviral vector pLV[Exp]-Puro-EF1A>dCas9/KRAB/MeCP2 and Mammalian Dual-gRNA Expression Lentiviral Vector with 2 synthetic sgRNA sequences (AAGGCATTACAGGTATGCAG, TATCAACCAAACGGCTTTGT), targeting glioma-specific cis-regulatory element or Control Vector with 2 sgRNA sequences (GTGTAGTTCGACCATTCGTG, GTTCAGGATCACGTTACCGC) were obtained from VectorBuilder (Chicago, IL, USA). GSCs and NHAs were seeded in 6-well plates and transfected with 500 ng of both vectors using Lipofectamine 2000 (Thermo Fisher Scientific) according to the manufacturer’s protocol. At 48 hours post-transfection, cells were harvested for ChIP-seq, RT-qPCR, 3C-qPCR, SPLASH-qPCR and WST-1 cell proliferation assays.

### Bulk Gro-Seq and data analysis

Nuclei were isolated following a previously described step-by-step fastGRO protocol (https://doi.org/10.17504/protocols.io.bbmgik3w). Briefly, 5 million cells were washed with ice-cold PBS twice and incubated in swelling buffer (10 mM Tris-HCl pH 7.5, 2 mM MgCl2, 3 mM CaCl2, and 2 U/mL Superase-In) for 5 minutes on ice. After a subsequent wash with swelling buffer supplemented with 10% glycerol, cells were treated with lysis buffer (10 mM Tris-HCl pH 7.5, 2 mM MgCl2, 3 mM CaCl2, 10% glycerol, 1% Igepal CA-630, and 2 U/mL Superase-In) to release the nuclei. The isolated nuclei were washed twice in lysis buffer and resuspended in freezing buffer (50 mM Tris-HCl pH 8.3, 40% glycerol, 5 mM MgCl2, and 0.1 mM EDTA) at a density of 20 million nuclei per 100 μL. Aliquots were flash-frozen on dry ice and stored at -80 C.

#### Normalized Nuclear Run-On

For the nuclear run-on (NRO) reaction, nuclei were thawed on ice and, where applicable, supplemented with spike-in nuclei. An equal volume of pre-warmed 2x NRO reaction buffer (10 mM Tris-HCl pH 8, 5 mM MgCl2, 300 mM KCl, 1 mM DTT, 500 μM ATP/GTP/4-thio-UTP, 2 μM CTP, 200 U/mL Superase-In, and 1% Sarkosyl) was added to nuclei. The reaction was incubated at 30 C for 7 minutes.

#### RNA Extraction and Fragmentation

RNA was extracted using TRIzol LS reagent (Invitrogen) according to the manufacturer’s instructions and recovered by ethanol precipitation. RNA concentration was determined via the Qubit High Sensitivity Assay (Invitrogen). RNA was fragmented using RNA Fragmentation Reagents (AM8740, Invitrogen). Fragmentation efficiency was verified using 1% agarose gel.

#### Biotinylation and Thiol-Specific Enrichment

Fragmented RNA was incubated in Biotinylation Solution (20 mM Tris pH 7.5, 2 mM EDTA, 40% DMF, and 200 μg/mL EZ-link HPDP-Biotin; Thermo Scientific) for 2 hours at 25 C (800 rpm) in the dark. Following ethanol precipitation, biotinylated RNA was resuspended in nuclease-free water and isolated using M280 Streptavidin Dynabeads (Invitrogen). Beads (100 μL per sample) were washed twice with two volumes of freshly prepared wash buffer (100 mM Tris pH 7.5, 10 mM EDTA, 1 M NaCl, and 0.1% v/v Tween-20), resuspended in one volume of wash buffer, and added to the RNA. After a 15-minute rotation at 4 C, the beads underwent three washes with a pre-warmed (65**°** C) wash buffer and three washes with room-temperature wash buffer. The 4-thio-UTP-incorporated RNA was eluted in 100 mM DTT and purified using the RNA Clean & Concentrator kit (Zymo Research) with an on-column DNase I digestion.

#### Library Preparation and Sequencing

Eluted RNA was quantified by Qubit and used for strand-specific library construction using the NEBNext Ultra II Directional RNA Library Prep kit (New England Biolabs). Libraries were sequenced on the Illumina NextSeq 2500 platform.

#### Data analysis

Sequencing reads were aligned to the human reference genome (hg38) using **STAR**^37^ **v2.5** in 2-pass mode. Reference indexes were constructed using the latest annotations from **Gencode Release 49**. The alignment was performed with the following parameters: -- quantMode TranscriptomeSAM, --outFilterMultimapNmax 10, --outFilterMismatchNmax 10, -- outFilterMismatchNoverLmax 0.3, --alignIntronMin 21, --alignIntronMax 0, -- alignMatesGapMax 0, --alignSJoverhangMin 5, and --twopassMode Basic. For the two-pass alignment, --twopass1readsN was set to 60,000,000 with a --sjdbOverhang of 100. Post-alignment, BAM files were filtered for mapping quality (q >= 10) using **SAMtools v1.13.7**. To account for experimental variation, BAM files were normalized using a scaling factor derived from the ratio of *Drosophila* spike-in reads to the total number of mapped reads. Finally, bigWig tracks were generated for visualization using **deepTools v3.5.0**.

### Bulk Circular Chromosome Conformation Capture with sequencing (4C-Seq), bulk 4C- qPCR and data analysis

The 4C libraries were prepared by the established protocol^38^ with minor changes. Briefly, cells were trypsinized, counted, and fixed at 10^6^ cells/ml with 1% formaldehyde (Sigma) in PBS for 15 min at RT. The fixation was quenched with cold glycine at a final concentration of 125 mM, the cells were washed with PBS and permeabilized on ice for 1 h with 10 mM Tris-HCl, pH 8, 100 mM NaCl, 0.1% NP-40, and protease inhibitors. Cell nuclei were resuspended in *Nlalll* restriction buffer to a concentration of 5×10^6^ nuclei/ml, further permeabilized in 0.4% SDS at 37°C, followed by the SDS neutralization with 2.6% Triton-X100 at 37°C. The nuclei were digested overnight with 300 U *Nlalll* at 37°C, and the enzyme was inactivated by incubation at 65°C for 30 min, followed by two washes and resuspension in T4 DNA ligase buffer. *In situ* ligation was performed in 3.5 ml T4 DNA ligase buffer with 100 U of T4 DNA ligase overnight at 16°C. The purified DNA was further digested with 5 U/μg of DpnII at 37°C overnight, then re-purified by phenol/chloroform extraction and isopropanol precipitation. The DNA was then circularized by ligation with 10 U/μg T4 DNA ligase under diluted conditions (5 ng/μl DNA). DNA was purified by reverse crosslinking with an overnight incubation at 65°C with proteinase K, followed by RNase A digestion for 30 min at 37°C, phenol/chloroform extraction, and isopropanol precipitation. 100 ng aliquots of ligated DNA were used as a template for PCR with bait-specific primers containing Illumina adapters. The products of the PCR reactions were pooled, the primers removed by washing with 1.8× AMPure XP beads, and then sequenced by HiSeq 2500 SE50 (Illumina). All experiments were performed using two biological replicates per condition. The primers are listed in Supplementary Table 3.

The first 20 nucleotides corresponding to the viewpoint-specific forward reading primer were trimmed from the raw sequencing reads, enabling mapping of the captured fragments. The reads were aligned with Bowtie2 (version 2.5.3) against the human reference genome (GRCh37 / hg19). The resulting Bam files were processed with functions from Bioconductor packages: r3Cseq^39^ (Bioconductor version 1.34.0). Briefly, the number of reads per restriction fragment was quantified and normalized (reads per million). The resulting bedgraph files were visualized using IGV web application^40^.

### Bulk 4C-qPCR

After 4C libraries were generated, DNA was amplified with primers listed in **Supplemental Table 3.** qRT-PCR was performed with SYBR GREEN chemistry and normalized to GAPDH loop for restriction and ligation efficiency. Relative number of DNA-DNA interactions normalized to GAPDH was calculated in two steps: 1) DDCt = (Ct IP) - (Ct GAPDH loop); 2) (2^-DDCt).

### Bulk SPLASH-qPCR

To profile RNA-RNA interactions, we adapted the SPLASH-seq^14^ protocol for quantitative PCR (qPCR) readouts.

**1) In vivo crosslinking and RNA extraction:** Cells were incubated with 200 μM EZ-Link Psoralen-PEG3-Biotin (Thermo Fisher) and 0.01% (w/v) digitonin at 37°C for 5 min, followed by irradiation with 365-nm UV light on ice for 20 min. Total RNA was extracted using TRIzol. **2) RNA fragmentation and Enrichment:** Total RNA (20 μg) was chemically fragmented at 95°C for 5 min (9 mM MgCl2, 225 mM KCl, 150 mM Tris-HCl, pH 8.3) and resolved on a 6% TBE-8M urea gel. The 90–110 nt fraction was excised and eluted overnight at 4 °C. Crosslinked RNAs (1.5 μg) were captured using 100 μL Dynabeads MyOne Streptavidin C1 (Life Technologies) in a formamide-containing hybridization/lysis buffer matrix at 37 °C for 30 min, followed by five washes with 2x SSC / 0.5% SDS. **3) Proximity ligation and reverse crosslinking:** Bead-bound RNA was phosphorylated using T4 PNK (NEB; 0.5 U, 37 °C for 4 h, followed by supplementation with 1 mM ATP for an additional 1 h). Interacting RNA strands were ligated using T4 RNA Ligase 1 (NEB; 2.5 U/μL) overnight at 16 °C. Chimeric RNA was eluted in SDS-containing proteinase K buffer at 95 °C for 10 min, purified using RNeasy Cleanup (Qiagen), and photo-reverse-crosslinked by irradiation with 254-nm UV light on ice for 5 min. **4) Adaptor ligation and cDNA synthesis:** Purified RNA was hybridized to 6 μM 3′ adaptors and ligated using T4 RNA Ligase 2 KQ (NEB) at 25 °C for 2.5 h. Products of 110–130 nt were size-selected by TBE-urea gel electrophoresis. Eluted RNA was reverse-transcribed using 208 nM RT primers and SuperScript III (Invitrogen) at 50 °C for 30 min. RNA templates were degraded with 100 mM NaOH at 98 °C for 20 min, and cDNAs of 200–220 nt were size-selected by gel electrophoresis. Purified cDNA was circularized using CircLigase II (Epicentre) and cleaned with a DNA Clean & Concentrator-5 kit (Zymo Research) before downstream qPCR analysis.

qRT-PCR was performed with SYBR GREEN chemistry using U4/U6 interaction primers as a normalization control. Relative RNA–RNA interaction abundance was calculated by first determining ΔΔCt = Ct(IP) − Ct(U4/U6), followed by calculating 2^−ΔΔCt. Primers are listed in **Supplemental Table 3**.

### Quantitative Real-Time PCR (qRT-PCR) and Data Analysis

Total RNA was isolated using the RNA Purification Kit (Norgen Biotek Corporation, #37500) according to the manufacturer’s protocol. For cDNA synthesis, 100 ng of total RNA was reverse transcribed using the iScript Reverse Transcription kit (Biorad, #1708841), followed by qPCR SYBR Green assays (Applied Biosystems, #4309155) with custom primers listed in **Supplemental Table 3**. Fold-change in gene expression between conditions was calculated by the 2^-ΔCt method.

### Combined DNA and RNA interactomics

#### Cell dissociation to single-cell suspension

Primary GBM tumor tissues were flash-frozen and stored at -80°C. Tissue was minced into ∼1 mm³ fragments and enzymatically dissociated in Hibernate-A medium (Thermo Fisher Scientific, Cat. No. A1247501) supplemented with papain (20 U/mL; Worthington Biochemical, Cat. No. LS003126), DNase I (0.005%; Sigma-Aldrich, Cat. No. DN25), and 1 mM L-cysteine at 37 °C for 30 min with gentle agitation. Digestion was terminated by adding Dulbecco’s modified Eagle medium/F12 (DMEM/F12; Thermo Fisher Scientific, Cat. No. 11320033) containing 10% fetal bovine serum (FBS; Gibco, Cat. No. 26140079). The suspension was gently triturated using fire-polished Pasteur pipettes, passed through a 40-µm cell strainer (Falcon, Cat. No. 352340), and centrifuged at 300 × g for 5 min. Cells were resuspended in PBS (Gibco, Cat. No. 10010023), and viability was assessed by trypan blue exclusion (Gibco, Cat. No. 15250061). Glioma stem cell lines were dissociated into single cells using the Neurocult Stem Cells Chemical Dissociation kit ( STEMCELL Technologies). Only single-cell suspensions with viability >85% were used for downstream analyses.

#### RNA-to-RNA and DNA-to-DNA crosslinking, fragmentation and DNA-to-DNA ligation

First, cells were incubated with 10 mL PBS containing 200 µM biotinylated psoralen and 0.01% (w/v) digitonin at 37 °C for 5 min, followed by irradiation with 365-nm UV light for 20 min on ice at a distance of 3 cm using a UV crosslinker (RNA-RNA crosslinking). Second, cells were fixed in 1% formaldehyde (Sigma-Aldrich) in PBS for 15 min at room temperature, and crosslinking was quenched by addition of cold glycine to a final concentration of 125 mM, followed by three washes with PBS (DNA-DNA crosslinking). Cells were then permeabilized on ice for 1 h in a buffer containing 10 mM Tris-HCl (pH 8.0), 100 mM NaCl, 0.1% NP-40, and protease inhibitors. Nuclei were resuspended in NlaIII restriction buffer at a concentration of 5 × 10⁶ nuclei/mL and further permeabilized with 0.4% SDS at 37 °C, followed by SDS neutralization with 2.6% Triton X-100 at 37 °C. Chromatin was digested overnight with 300 U NlaIII at 37 °C, followed by heat inactivation at 65 °C for 10 min, which simultaneously induced RNA fragmentation. After two washes, nuclei were resuspended in T4 DNA ligase buffer, and *in situ* DNA ligation was carried out in a 3.5 mL ligation mix containing 100 U T4 DNA ligase for 4 h at 16 °C.

Nuclei were distributed into 96-well plates by limiting dilution. Nuclear membranes were lysed and chromatin was reverse-crosslinked by incubation with Proteinase K for 4 h at 65 °C followed by 15 min at 95 °C to deactivate Proteinase K. Biotinylated RNA complexes were captured using streptavidin magnetic beads and transferred to fresh wells. DNA interactomic library preparation was performed using a two-step PCR approach. Locus-specific amplification was conducted using custom primers (**Supplemental Table 3, PCR Round 1**) containing Nextera-compatible universal adapter handles. Multiplexing was achieved in a second PCR step using **Illumina DNA/RNA Unique Dual Indexes (UDI) Set A barcodes** (**Supplemental Table 3, PCR Round 2**). Libraries were sequenced on an Illumina HiSeq 2500 at 300-cycle paired-end sequencing (2×150bp). Fastq forward and reverse reads for each sample were merged using FLASH^13^ (version 1.2.11) and deposited in the GEO Database.

#### RNA–RNA proximity ligation and library preparation

For 3′ RNA end repair, bead-enriched RNA–RNA crosslinked material was washed in ice-cold T4 polynucleotide kinase (PNK) buffer and treated with 0.5 U T4 PNK (NEB) in an 80-µL reaction at 37 °C for 4 h. Subsequently, fresh ATP (1 mM final) and an additional 0.5 U T4 PNK were added in a total volume of 100 µL and incubated for 1 h at 37 °C to repair 5′ RNA ends, followed by the buffer removal. RNA–RNA chimeras were generated by ligation in T4 RNA ligase buffer (NEB) supplemented with 1 mM ATP, 0.5 U/µL RNase inhibitor, and 2.5 U/µL T4 RNA ligase 1 (NEB) at 37 °C for 1 h with continuous agitation. Reverse crosslinking was completed by UV irradiation at 254 nm for 5 min on ice. RNA on streptavidine beads was washed once with 1x scRT Buffer (Takara, #634444). RT and subsequent steps were performed using the SMART-Seq® Stranded Kit (Takara, #634444), following the manufacturer’s instructions. Libraries were sequenced on the Illumina HiSeq 2500 using 300-cycle paired-end sequencing (2×150bp). Fastq forward and reverse reads for each sample were merged using FLASH^41^ (version 1.2.11) and deposited in GEO.

### Bioinformatic analyses of combined DNA and RNA interactomics

The DNA-DNA interactomics wet-lab protocol consists of two primary enzymatic stages: (1) *in situ* genomic DNA digestion using the NlaIII restriction endonuclease, targeting the 5’-CATG-3’ recognition sequence, and (2) T4 DNA ligase-mediated proximity ligation, which facilitates the formation of chimeric DNA-DNA junctions from spatially proximal fragments. Reads were splitted by Cell barcodes (**Supplemental Table 3**). Chimeric DNA–DNA reads consist of a viewpoint-specific sequence followed by one or more genomic fragments separated by CATG sites. Reads were split at CATG motifs using a custom awk script (Mendeley Data, doi: 10.17632/7zzry8fr5f.1), deduplicated and individual segments were aligned to the human hg38 genome using Bowtie2 (default parameters). Parsed results, including cell ID, viewpoint, segments, and their genomic coordinates are provided in GSE336543 (xlsx file).

RNA-RNA interacting fragments were extracted, processed, and quantified as follows: FLASH-merged (version 1.2.11) forward (Read 1) reads, which match the antisense sequence of the input RNA were extended. Duplicate reads were collapsed into a fasta file, with duplicate counts per read calculated using fastx_toolkit (version 0.0.14). Chimeric reads were aligned to a reference fasta file containing full transcript sequences with introns (Gencode V50 fasta extended with LNCipedia v5.2) using BWA-MEM (v0.7.17) in local alignment mode considered transcript strand only (antisense to Read 1 extended). Alignment parameters included: Seed length for the first mapping iteration = 16, for the second mapping iteration = 18; Minimum alignment score for the first iteration = 20, for the second iteration = 18; Matching score for the first iteration = 1, for the second iteration = 1; Mismatch penalty for the first iteration = 4, for the second iteration = 6; Gap opening penalty for the first iteration = 6, for the second iteration = 100; Gap extension penalty for both iterations = 1; Maximum multi-hit reads allowed: 50 for the first iteration, 100 for the second iteration. Pairs of genes from each chimeric read were identified, and the number of reads for each gene pair was quantified with ChiRA^42^ (v1.4.3).

To evaluate the enrichment of RNA–RNA interaction pairs above random expectation, we implemented a probabilistic framework in R (v3.6.2; base stats package) using a one-sided exact binomial test. For each RNA pair (A, B), the null probability of interaction (P_exp) was defined under a random collision model as the product of their relative abundances, calculated as P_exp = (n_A/N_total) x (n_B/N_total), where n represents individual transcript abundance and N_total is the aggregate abundance across the dataset. Observed interaction counts (k) were tested against this null expectation, with the total number of detected interaction events in the experiment serving as the number of trials. We reported the expected interaction probability, exact p-value, and 95% confidence intervals for the binomial proportion for all tested pairs. To account for multiple hypothesis testing, p-values were adjusted using the Benjamini–Hochberg procedure to control the false discovery rate (FDR), and interactions with an FDR-adjusted p-value < 0.05 were considered significantly enriched and are shown on all Figures of this article (Mendeley Data, doi: 10.17632/7zzry8fr5f.1). Custom awk scripts were used to generate a count table (rows: gene pairs, columns: cells), which was further processed using functions from Seurat^43^ (version 5) to scale, normalize the data, define and visualize cell clusters.

### Combined ASO barcodes-seq and scRNA-seq

Human organoids were treated with barcoded antisense oligonucleotides (ASOs) in LNP formulations. Following a 5-day incubation period, the organoids were mechanically and chemically dissociated into a single-cell suspension using the NeuroCult Chemical Dissociation Kit for Mouse CNS Stem Cells (STEMCELL Technologies) according to the manufacturer’s protocol.

Cells were exposed to 365 nm UV for 5 minutes to cleave iPC Linker connecting ASO to its unique barcode (see ASO design). Dead cells were separated with EasySep™ Dead Cell Removal (Annexin V) Kit (Catalog #17899). Both dead and viable single cells were deposited into individual wells of a 96-well plate, having a PCR-compatible Proteinase K-based lysis buffer (10 mM Tris-HCl (pH 8.0), 50 mM KCl, 2.5 mM MgCl2, 0.45% NP-40, 0.45% Tween-20, Proteinase K (100 µg/mL)), followed by 55°C, 30 min incubation to disrupt both cell and nuclear membranes and then heat-inactivation of the Proteinase K at 95°C for 10 minutes. Long polyadenylated (polyA) RNA (mRNA and lncRNA) was captured using Dynabeads Oligo(dT)25 magnetic beads (Thermo Fisher Scientific, Cat. No. 61002) and transferred to a separate 96-well processing plate for transcriptome library preparation.

ASO Barcodes (DNA) from different wells were amplified for Illumina sequencing with the combination of cell barcodes unique for each well (**Illumina DNA/RNA Unique Dual Indexes (UDI)**, listed at **Supplemental Table 3):**

Primer_1: 5’-AATGATACGGCGACCACCGAGATCTACAC**[i5_Index**]

ACACTCTTTCCCTACACGACGCTCTTCCGATCT - 3’

Primer_2: 5’-CAAGCAGAAGACGGCATACGAGAT**[i7_Index]**

GTGACTGGAGTTCAGACGTGTGCTCTTCCGATCT - 3’

The resulting amplicon library architecture was arranged as follows:

P5 Flow Cell Anchor — [i5 Index] — Read 1 Primer — [Barcode] — Read 2 Primer — [i7 Index] — P7 Flow Cell Anchor

Single-end sequencing (150 bp) was performed using an Illumina HiSeq 2500. Resulting fastq files were demultiplexed based on cell barcodes, followed by the quantification of unique 25 nt long random sequences [Barcode] for each cell.

Reverse transcription (RT), template switching, and secondary cDNA amplification of the isolated polyA-selected mRNA fractions were conducted using the SMART-Seq Stranded Kit (Takara Bio, Cat. No. 634444) in strict accordance with instructions. The completed multiplexed single-cell RNA-seq cDNA libraries were sequenced on an Illumina HiSeq 2500 system using a 300-cycle paired-end (2×150 bp) configuration. Forward and Reverse reads of each fastq file were merged with FLASH (v1.2.11, Minimum overlap=1). Resulting reads were aligned to GRCh38/hg38 genome assembly with Hisat2^44^ (v2.2.1; default parameters). Number of reads in genes were quantified with featureCounts^45^ for each cell (v2.1.1; using Gencode V50) and combined to a single table, compatible with Seurat (version 5) to scale, normalize the data, define and visualize cell clusters.

### Sequential single-molecule FISH (seq-smFISH)

#### Cell culture and tissue preparation

Cells were cultured under standard conditions and plated onto poly-L-lysine-coated glass coverslips at a density of 20,000 cells/cm². Frozen tissue was sectioned into 10-µm slices using a cryostat and mounted onto glass slides. At the time of fixation, cells were washed twice with phosphate-buffered saline (PBS) and fixed in 4% paraformaldehyde (Electron Microscopy Sciences, #15710) in PBS for 10 min at room temperature. Following fixation, samples were washed three times with PBS and permeabilized in 70% ethanol at 4°C overnight.

#### Probe design

Primary probe sets targeting five lncRNAs (HOTAIRM1, HOXB-AS1, HOTAIR, HOXD-AS2, HOXDeRNA) were designed using Stellaris Probe Designer (version 4.2) with following parameters: Organism - Human, Masking Level - 5, Max Number of Probes - 48, Oligo Length - 20, Min Spacing Length - 2 nt. Each probe set consisted of 30–48 DNA oligonucleotides tiled along the first intron of each lncRNAs. Oligonucleotides were synthesized by Biosearch Technologies and are listed at **Supplemental Table 4**.

#### Sequential probe hybridization

Sequential detection of the five lncRNAs was performed following Stellaris RNA FISH protocols for both tissues and adherent cell lines (Biosearch Technologies, https://oligos.biosearchtech.com). For each round, samples were incubated with fluorescent probes. Excess probes were removed by two PBS washes. After completion of all washes, nuclei were stained with DAPI (1 µg/mL in PBS, 5 min). Coverslips were mounted in an antifade mounting medium (ProLong Gold, #P36930). The images were acquired using a Leica DMi8 fluorescence microscope with a 63×NA 1.4 oil objective. After imaging, fluorophores were stripped using 0.5U/µL DNAse I (Roche). To inhibit DNase I, cells were washed 2X with 30% Formamide, 0.1% Trition-X 100, 2X SSC prior to the next hybridization round.

#### Measurement of distances between lncRNAs

Single-molecule lncRNA foci were identified using semi-automated segmentation in ImageJ (v1.54g; NIH). For tumor samples, color thresholding was performed in HSB color space independently for each channel. For time course experiments, image brightness was adjusted to maximize signal-to-noise ratio and suppress background fluorescence, and binary thresholding was applied independently to each grayscale channel to isolate fluorescent foci. Spot centroid coordinates (X, Y) were extracted using the

Analyze Particles function (size filter: 2–40 pixels2; edge particles excluded). Pixel coordinates were converted to micrometers using a calibration factor of 0.102 μm/pixel. Pairwise inter-lncRNA distances were computed using custom Python scripts (Python 3.12; pandas v2.1.4, NumPy v1.26.4, SciPy v1.11.4, Matplotlib v3.8.0). For all ten pairwise combinations of the five lncRNA species, Euclidean distances were calculated between spot centroids using scipy.spatial.distance.cdist. For each lncRNA A spot, the minimum distance to any lncRNA B spot was recorded to quantify nearest-neighbour associations. Analysis was restricted to intra-nuclear measurements by applying a 5.0 μm maximum distance threshold, approximating nuclear diameter. All pairwise combinations were pooled within each patient sample (Figure 5F) or within each timepoint (Figure 6G).

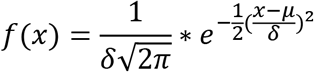

where μ is the mean distance and σ is the standard deviation, estimated via maximum likelihood using scipy.stats.norm. Per-patient statistics (mean µ, standard deviation σ, standard error s.e.m. = σ/√n, where n is the number of distance measurements) were computed independently. Population-level statistics were derived from per-patient means. Inter-patient variability was quantified by coefficient of variation (CV = 100 × σ_patients/µ_overall). Visualization was performed using Matplotlib. All custom scripts sufficient to reproduce the analysis were deposited to Github (https://github.com/kinseyam/lncRNA_distance-intensity_quantification).

#### Quantification of lncRNA Fluorescence Intensity

Single-molecule lncRNA foci were identified independently in each channel using semi-automated segmentation in ImageJ (v1.54g; NIH). Binary thresholding was applied to each grayscale channel separately to isolate fluorescent puncta. Mean fluorescence intensity values were extracted for each detected spot using the Analyze Particles function (size filter: 2–40 pixels²; measurements: mean gray value). Intensity distributions were analyzed using custom Python scripts (Python 3.12; pandas v2.1.4, NumPy v1.26.4, SciPy v1.11.4, Matplotlib v3.8.0). All lncRNA species were pooled within each timepoint for statistical modelling. Per-timepoint statistics (mean µ, standard deviation σ, standard error s.e.m. = σ/√n, where n is the number of detected lncRNA foci) were computed across all channels combined. Intensity distributions were modeled as Gaussian (normal) probability density functions. For each timepoint, the probability density f(I) at intensity I was calculated as:

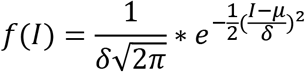

where μ is the mean intensity and σ is the standard deviation, estimated via maximum likelihood using scipy.stats.norm.pdf. The resulting probability density values (y-axis) represent the relative likelihood of observing each intensity value, normalized such that the total area under the curve equals 1. This normalization allows direct comparison between timepoints despite different sample sizes. Data visualization was performed using Matplotlib. All custom scripts sufficient to reproduce the analysis were deposited to Github (https://github.com/kinseyam/lncRNA_distance-intensity_quantification).

### Alternative Splicing Analysis

#### RNA-sequencing Data Acquisition and Preprocessing

Publicly available total RNA-sequencing datasets were obtained from the Gene Expression Omnibus (GEO) database for: GBM primary tumors, glioma stem cells, neural stem cells (GSE119834); astrocytes, oligodendrocytes, OPCs, neurons, myeloid cells, along with whole cortex samples (GSE73721, GSE239662, GSE99349).

#### Splicing Quantification with Whippet Pipeline

Alternative splicing analysis was performed using the Whippet computational pipeline^20^ (v1.6), an alignment-free method optimized for accurate and efficient splicing quantification. A transcriptome index was constructed using the comprehensive GENCODE v50 human genome annotation (GRCh38/hg38 assembly). Individual sample quantification was conducted with ‘whippet-quant’, generating transcript-specific read counts and splicing graphs.

#### Differential Splicing Analysis

Pairwise differential splicing comparisons between GBM or GSC samples and each reference brain cell type were performed using ‘whippet-delt’. This analysis calculates ΔΨ values (change in percent spliced in) and associated Bayesian posterior probabilities for each splicing event. ΔΨ represents the difference in isoform inclusion rates between conditions, with positive values indicating increased inclusion in GBM or GSC and negative values indicating decreased inclusion. GBM and GSC results were intersected for the common list of genes.

#### Statistical Filtering and Event Annotation

Differential splicing events were filtered using stringent significance thresholds: |ΔΨ| ≥ 0.50 (moderate to large effect size) and posterior probability ≥ 0.90 (high confidence). Each significant event was assigned a unique EventID composed of the gene identifier, splicing event type (Core exon [CE], alternative acceptor [AA], alternative donor [AD], retained intron [RI], tandem transcription start site [TS] or Tandem alternative polyadenylation site [TE], Alternative First exon [AF], Alternative Last Exon [AL]), and genomic coordinates. Ensembl gene identifiers were mapped to official HUGO gene symbols.

### Integration of HiC with RNA-Seq

Topologically Associating Domains (TADs) were identified across five human cell types using Hi-C datasets sourced from the ENCODE Consortium: astrocytes (ENCSR011GNI), macrophages (ENCSR236EYO), neurons (ENCSR228TUX), endothelial cell (ENCSR507AHE), and glioma stem cell (3 glioma stem cells from Johnston 2019^18^, https://wangftp.wustl.edu/hubs/johnston_gallo/), using the hicFindTADs function from HiCExplorer with parameters:

Matrix to compute on: **corrected_contact_matrix_large.h5 ;** Minimum window length (in bp) to be considered to the left and to the right of each Hi-C bin: **30000 ;** Maximum window length (in bp) to be considered to the left and to the right of each Hi-C bin: **100000 ;** Step size when moving from minDepth to maxDepth: **10000 ;** Multiple Testing Corrections: **False discovery rate;** q-value: **0.05 ;** Minimum threshold of the difference between the TAD-separation score of a putative boundary and the mean of the TAD-sep score of surrounding bins: **0.001.**

RNA-seq datasets for **neurons** (GSE99349), **oligodendrocytes/OPCs** (GSE239662), **astrocytes** (GSE73721), **endothelial cells** (GSE233210), and **macrophages** (GSE135491), were retrieved from GEO database. Reads were aligned to the **GRCh38 (hg38)** assembly using **HISAT2 (v 2.2.0)** with defaults, followed by gene quantification with **featureCounts (v2.0.6)**. Genes were assigned to their respective TAD coordinates. To identify the most highly correlated transcript pairs within the same TAD domain, **Pearson correlation coefficients** were calculated for all intra-TAD gene pairs across each cell type (Mendeley Data, doi: 10.17632/7zzry8fr5f.1). Interacting RNA segments from RNA SCIENCE were annotated to respective genes and respective TAD domains for **Figure 1G**. A list of gene pairs with the highest correlations was intersected with the list of interacting genes from RNA SCIENCE data and presented in **Figure 1H**.

## Data availability

Sequencing data generated in this study have been deposited at GEO: RNA SCIENCE (GSE337464), DNA SCIENCE (GSE336543), scRNA-seq on organoids (GSE337246), scBarcode-seq on organoids (GSE337339), ChIP-Seq (GSE337829), GRO-seq (GSE336545), 4C-seq (GSE336541). Microscopy data were deposited to Mendeley with doi:10.17632/7zzry8fr5f.1. The data are publicly available as of the date of publication.

## Code availability

All custom scripts sufficient to reproduce **Sequential single-molecule FISH (seq-smFISH)** imaging analysis were deposited to Github (https://github.com/kinseyam/lncRNA_distance-intensity_quantification). Custom R and awk scripts were deposited to: Mendeley Data, doi: 10.17632/7zzry8fr5f.1

## Contributions

AMK and ED conceived and designed the study; ED performed most experiments, data analysis, and visualization; YZ, HM, AJ, AK, ZZ and AEK assisted with experiments; YZ made LNP formulations; AK analyzed and visualized smFISH data; HM and AJ performed organoid generation and immunostaining; CEB provided resources, supervision, and funding acquisition for the organoid generation and immunostaining experiments; AEK contributed to data analysis; AMK supervised the work. ED and AMK wrote the manuscript and all authors revised and approved the manuscript.

## Corresponding author

Anna M. Krichevsky,

Evgeny Deforzh

## Ethics declarations

## Materials & Correspondence

Further information and requests for resources and reagents should be directed to the lead contact, Anna M. Krichevsky.

---

**Supplemental Figure 1.**
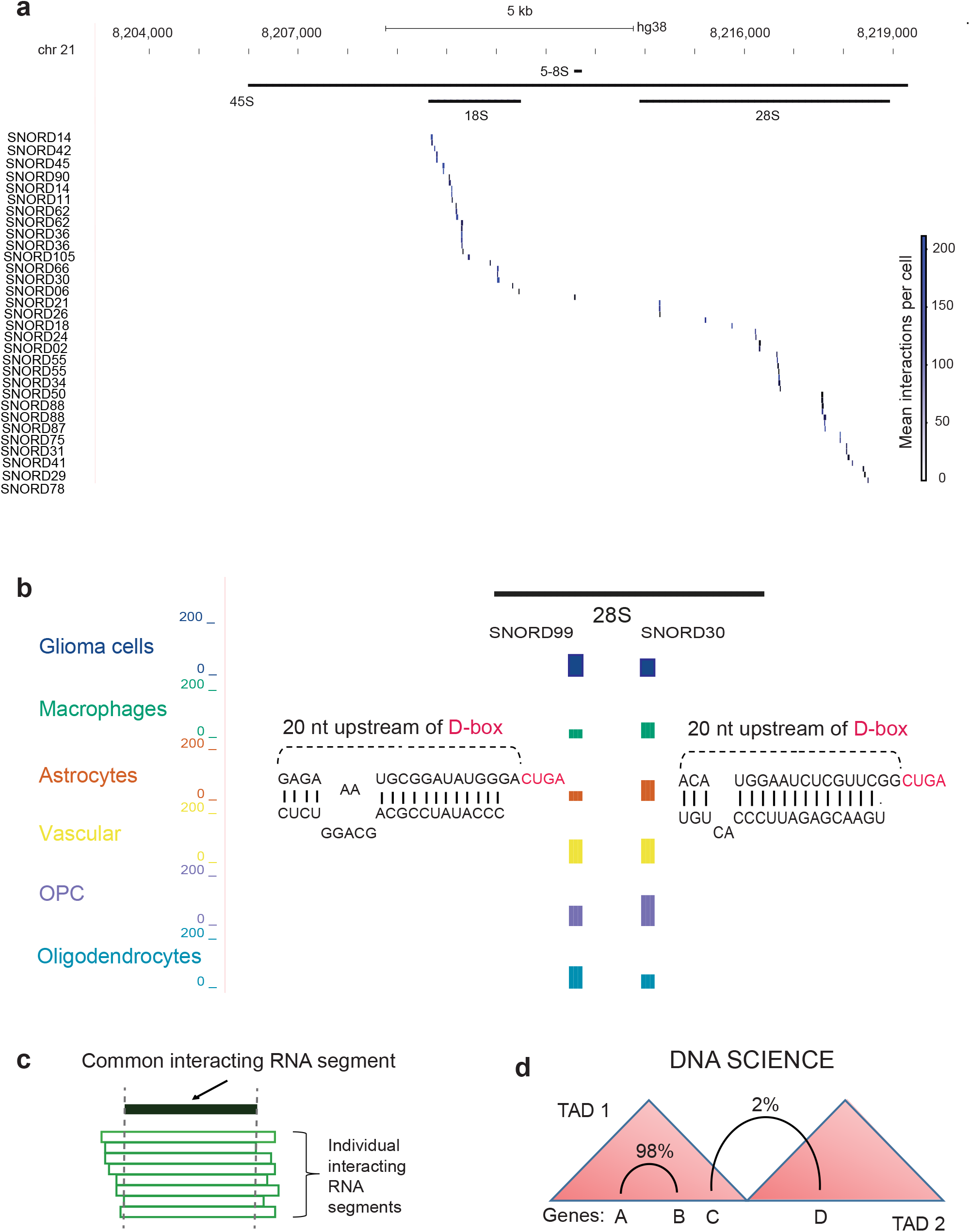
RNA and DNA SCIENCE interactomic profiling in primary glioma. Related to Figure 1. A. Canonical interactions between 45S pre-rRNA (top, black) and C/D box snoRNAs (SNORDs, blue) detected by RNA SCIENCE. Blue color gradient indicates mean interactions per cell. B. Cell-type-specific profiling of representative SNORD interactions aligned to 28S rRNA (black). Segments are color-coded by cell type, with bar height indicating mean interactions per cell. C. Individual interacting RNA segments (RNA SCIENCE) were merged to the Common interacting segment. D. Spatial distribution of interacting pairs of DNA segments relative to Topologically Associating Domain (TAD) boundaries. Integration of Hi-C and DNA SCIENCE datasets shows 98% of DNA interactions are intra-TAD constrained.

**Supplemental Figure 2.**
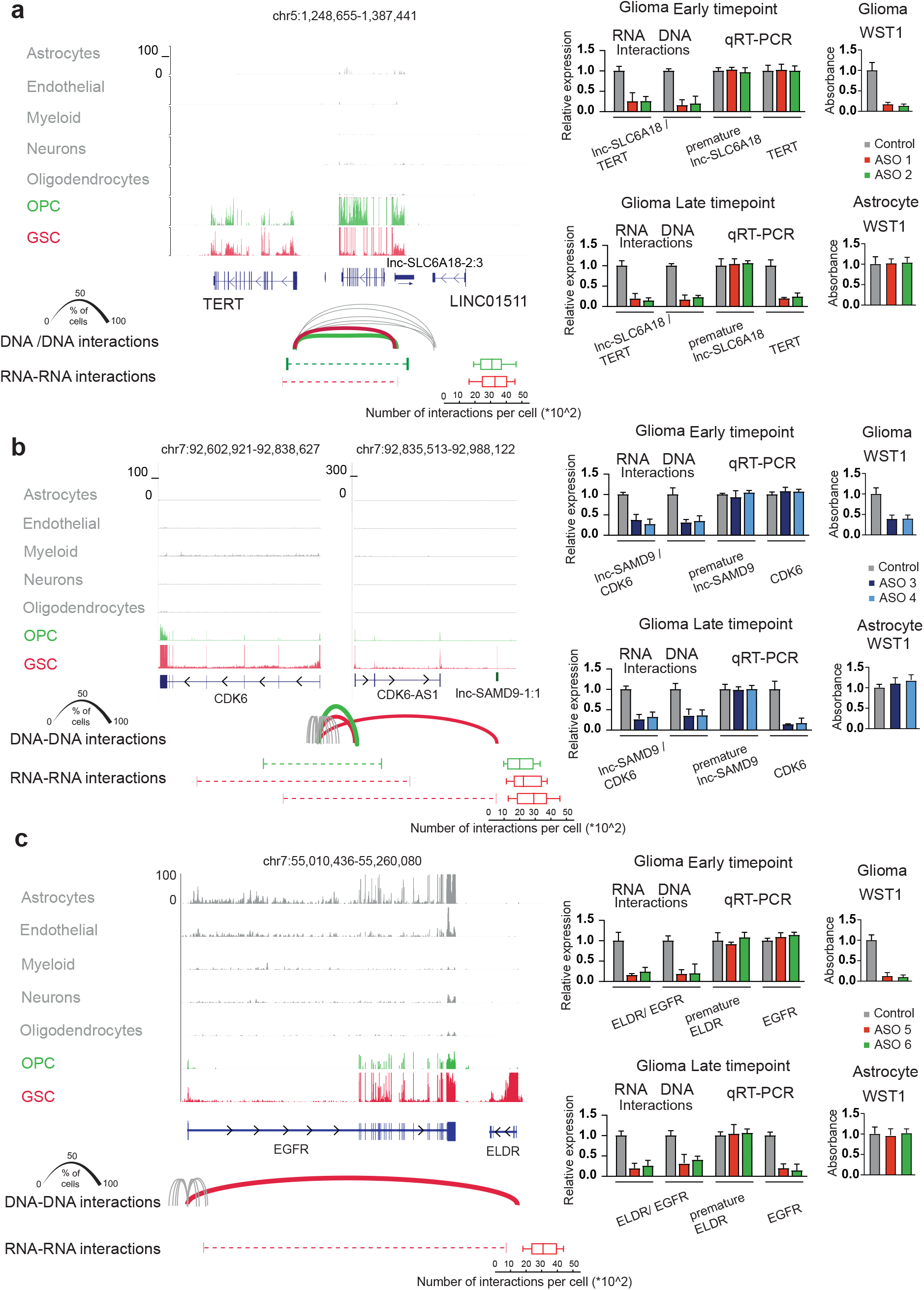
Targeted disruption of the cell-specific RNA and DNA interactomes of TERT, CDK6 and EGFR glioma oncogenes with ASOs. Related to Figure 2. A, B, C (Left). Total RNA-Seq visualized for corresponding glioma oncogene and surrounding lncRNAs in 7 cell types, followed by DNA-DNA (DNA SCIENCE; arcs) and RNA-RNA (RNA SCIENCE; bars connected with dotted lines) interactions. Arc width (DNA SCIENCE) represents the percentage of cells with a corresponding DNA-DNA interaction. Boxplots show the frequency of RNA SCIENCE contacts per cell between common interacting segments. A, B, C (Right). Treatment of glioma stem cells or astrocytes with two 2 ASOs disrupts cell type-specific premature RNA interactions (SPLASH-qPCR; n=3, mean with SD) and DNA loops (3C-qPCR; n=3, mean with SD) between corresponding glioma oncogene and cell-type specific lncRNA, leading to decreased oncogene expression (qRT-PCR, mean with SD, n=3) and glioma cell death (WST1, mean with SD, n=3). Astrocytes (WST1, mean with SD, n=3) remain intact. Early-timepoint analysis reveals that ASO treatment successfully dissociates RNA and DNA interactions in glioma prior to any measurable reduction in steady-state transcript levels.

**Supplemental Figure 3.**
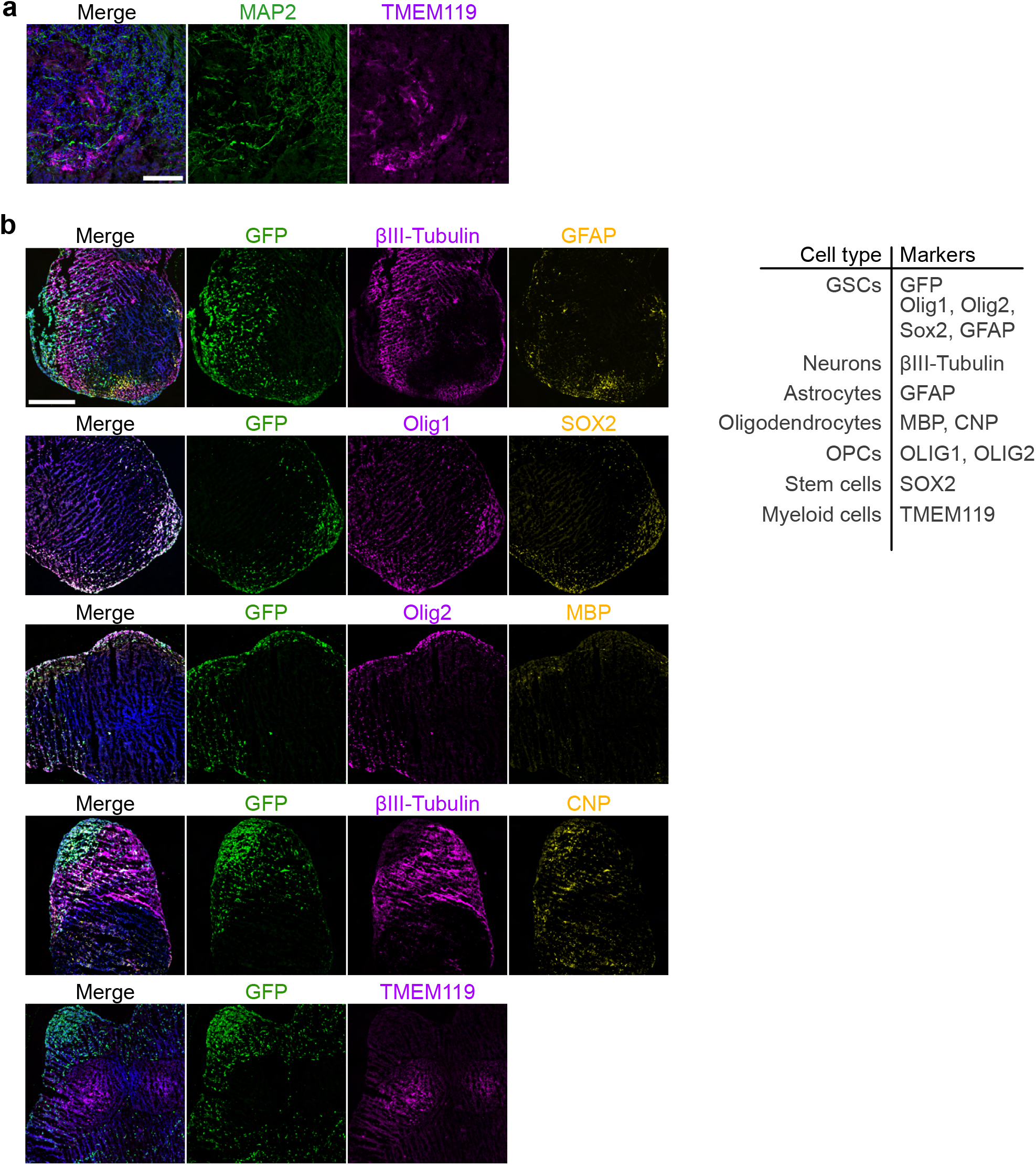
Characterization and GSC engraftment of cerebral organoid models. A. Representative immunofluorescence images of mature organoids (>70 days in culture) stained for TMEM119 (myeloid/microglial marker) and MAP2 (neuronal marker) to confirm myeloid cell differentiation. Scale bar = 100 µm. B. Representative immunostaining of GSC-engrafted organoids (day 95) showing GSCs (GFP) alongside lineage-specific markers: neurons (βIII-Tubulin), astrocytes (GFAP), oligodendrocytes (MBP, CNP), progenitor cells (SOX2), and microglia-like cells (TMEM119). Scale bar = 500 µm.

**Supplemental Figure 4.**
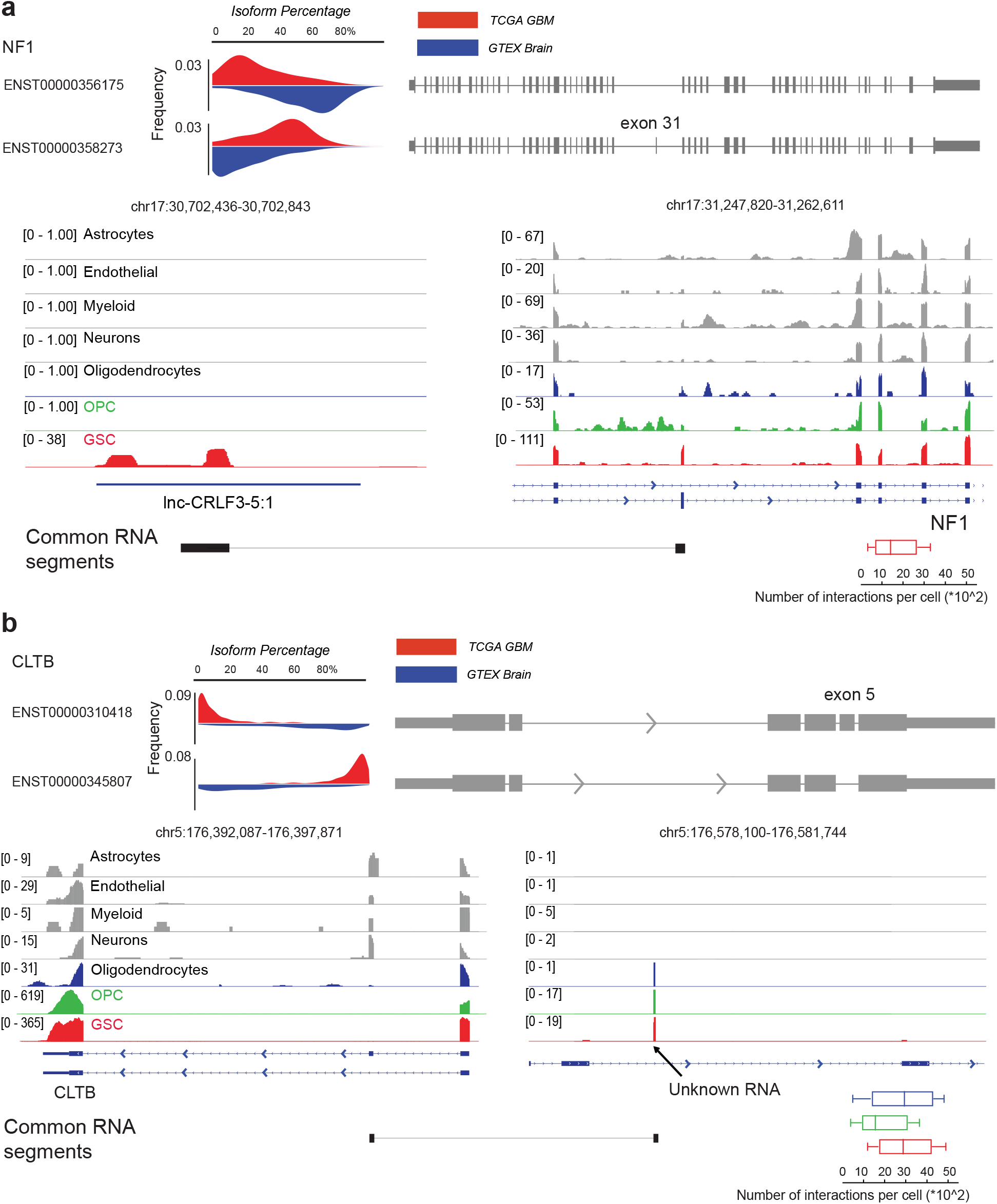
SCIENCE-Seq correlates with cancer-specific alternative splicing in glioma primary tumors. Related to Figure 4. A, B (Top) Isoform switching of NF1 or CLTB in glioma. Comparative analysis of NF1 or CLTB isoform percentage in glioma samples (TCGA, n=170) versus normal brain tissue (GTEx, n=870). Schematic representations of the corresponding transcript models are provided on the right. A, B (Bottom) Total Bulk RNA-Seq in different cell types showing cell type-specific lncRNA expression (top) followed by common RNA interacting segments measured by SCIENCE-Seq (bottom). Boxplots (bottom, right) quantify the frequency of each RNA–RNA interaction per cell.

**Supplemental Figure 5.**
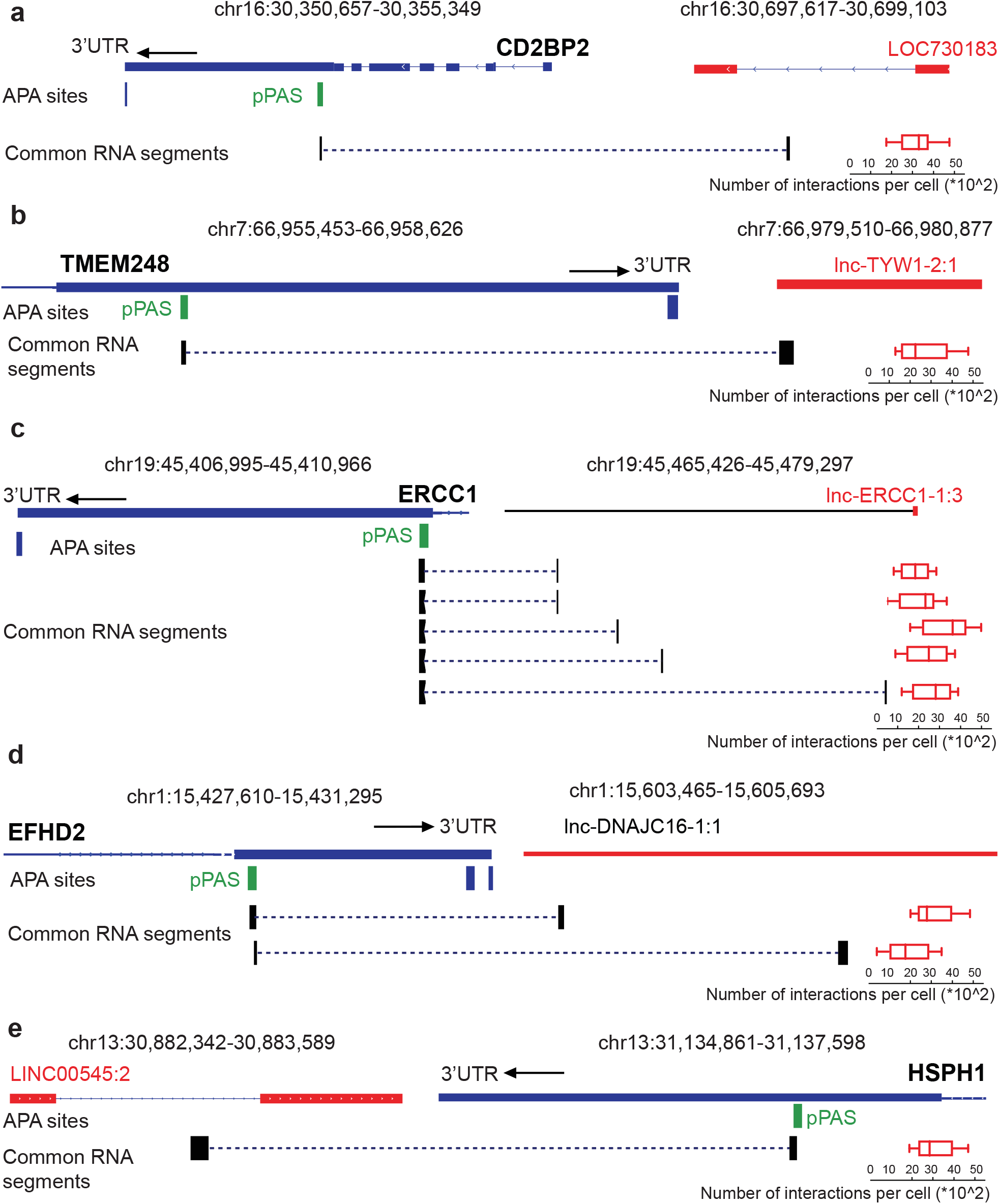
RNA SCIENCE detects lncRNA-pPAS interactions within glioma oncogene 3′UTRs. Related to Figure 5. (A–E) Proximal polyadenylation sites (pPAS, green) within the 3′UTRs of oncogenes (CD2BP2, TMEM248, ERCC1, EFHD2, and HSPS1) interacting with glioma-specific lncRNAs (red). Pairs of common interacting RNA segments are shown in black, with corresponding quantification per cell on the right (boxplots). Arrows indicate the directionality of the transcript (sense or antisense to forward DNA strand).

